# Molecular Determinants of Sequence-Dependent N-glycosylation by the Human Oligosaccharyltransferase

**DOI:** 10.64898/2026.09.03.749169

**Authors:** Milorad Andjelkovic, Juan Francisco Belén-Aguilar, Diego Belda, José Javier Ruiz-Pernía, Iñaki Tuñón, Ismael Mingarro

## Abstract

Protein N-glycosylation is catalysed by the oligosaccharyltransferase (OST) complex, yet how local sequon composition controls substrate selection by human OST remains poorly understood. Here we combine atomistic molecular dynamics, alchemical free-energy calculations, *in cellulo* glycosylation assays and human glycoproteome analysis to define the molecular basis of sequence-dependent N-glycosylation. We built, to our knowledge, the first atomistic model of human OST with both the lipid-linked oligosaccharide donor and an acceptor peptide simultaneously accommodated in a catalytically competent complex. Simulations show that conserved STT3 motifs stabilize the acceptor Asn and explain how substitutions at positions 0, +1 and +2 alter binding and productive geometry. Cellular assays in HEK 293T cells validate the predicted loss of glycosylation for N_0_Q sequon, +1 Pro and the +2 hierarchy Thr > Ser >> Cys. Analysis of 14,800 human glycosites confirms these rules at proteome scale.

## Introduction

Protein N-glycosylation is a near-universal co- and post-translational modification highly conserved across all domains of life.^1–3^ This process plays a crucial role in protein folding, maturation, quality control, trafficking, antigenicity, and many other biological processes.^4–6^ N-glycosylation specially affects membrane and secretory proteins and it is normally co-translationally catalysed by the oligosaccharyl-transferase (OST), a multi-transmembrane enzyme complex located in the endoplasmic reticulum (ER) and associated with the Sec61 translocon.^7,8^ During the process, the OST catalytic subunit STT3 mediates the transfer of a pre-assembled oligosaccharide (Glc_3_Man_9_GlcNAc_2_) from a dolichol pyrophosphate lipid carrier onto the side-chain amide of a target asparagine within the consensus sequon N-X-S/T, where “X” can be any amino acid except proline.^9–11^ This peptide specificity by OST is remarkably maintained in eukaryotes,^12,13^ archaea^14^ and bacteria,^15^ underscoring the evolutionary robustness of this motif for efficient N-glycosylation.

In humans, more than 7,000 proteins have been identified as N-glycosylated, based on current glycoproteomics data,^16^ reflecting the pivotal involvement of the OST complex in cellular physiology. Mammals express two OST isoforms; co-translational OST-A, which contains STT3A and scans nascent chains emerging from the ribosome, and post-translational OST-B, containing STT3B and acting as a proofreading mechanism to glycosylate sequons that OST-A may miss (e.g., in partially folded substrates or sequons close to C-terminal regions).^16–19^ Interestingly, in human cells OST-A constitutes the predominant oligosaccharyltransferase, while both isoforms can also compensate one another to secure efficient glycosylation.^20^ Recent high-resolution cryo-electron microscopy structures of human OST complexes have provided important insights into its architecture and offer a framework for probing key mechanistic aspects, including the roles of specific residues in catalysis, metal ion coordination, substrate docking, the contribution of accessory regions and how sequon context modulates glycosylation at the atomic level.^21,22^ These aspects comprise an important point in the field of biochemistry and biomedicine, since N-glycosylation defects are directly linked to human diseases,^23^ underscoring the importance of dissecting the chemical and molecular mechanisms that govern this essential modification.

In the canonical substrate sequon for N-glycosylation, the acceptor asparagine residue is defined as position “0”, residue Asn(0). Extensive previous works have shown that residues flanking the N-X-S/T motif strongly influence glycosylation efficiency.^11,24–28^ In particular, the residue at +2 is critical, with threonine generally yielding higher glycosylation rates than serine, a trend consistently observed across multiple studies.^11,24,25,29^ Remarkably, cysteine at the +2 position can create a non-canonical N-X-C sequon, suggesting that, under certain contexts, OST can act beyond the classical N-X-S/T consensus peptide.^30,31^ In contrast, there is general agreement that the presence of proline at +1 precludes glycosylation, likely due to its conformational rigidity disrupting recognition by OST.^11,27,32,33^ Together with the contributions of residues preceding the acceptor sequon (-2 and -1) these observations highlight that N-glycosylation efficiency emerges from a broad sequence context.^26,27,28^ . In addition, aromatic and bulky residues (such as Trp and Phe), as well as sulphur-containing residues (Cys and Met) in position +1, have been reported to strongly influence glycosylation outcomes as well.^11^ Nevertheless, most of these findings derive from *in vitro* studies such as microsomal assays, which, despite their sensitivity and reliable quantification, cannot fully capture the complexity present in living cells. To overcome these limitations, experiments in mammalian cells are essential to validate whether sequence-dependent effects observed *in vitro* translate into physiological contexts. At the same time, computational approaches provide a unique opportunity to bridge these two levels of analysis. Molecular dynamics (MD) simulations could enable exploration of the catalytic subunit STT3 and its interactions with substrates at atomic resolution, offering mechanistic insights inaccessible to static cryo-EM structures or bulk biochemical assays, as recently explored in the context of the N-glycosylation by the yeast OST.^34^ By probing transient states, substrate alignment and residue-specific contributions, MD can test and refine structural models. Integration with *in cellulo* validation, as made in this work, provides a coherent molecular framework to fully understand the sequence-dependent N-glycosylation efficiency from an atomic perspective, explicitly considering the contribution of individual residues to substrate binding.^35,36^

Herein, we report a combined approach, computational and cellular, to dissect the mechanism of N-glycosylation focusing on the chemistry of the process. Using the cryo-EM structure of the human OST (PDB 6S7O) complemented with AlphaFold^37^ predictions to resolve flexible loops and parametrizing the lipid-linked oligosaccharide (LLO) from the yeast OST structure (PDB 8AGC), we generated an atomistic model of the active site and proposed roles for key residues in metal coordination and catalysis. Free Energy simulations with a model substrate peptide sequence (^Nt^-A-Y-A-N-A-T-S-A-A-^Ct^) allowed us to quantify how substitutions at positions 0, +1, and +2 impact substrate binding. In parallel, we implemented a cellular assay based on the bacterial leader peptidase (Lep),^8,26,38,39^ as a membrane protein glycosylation reporter expressed in HEK 293T cells, enabling direct validation of the sequence-dependent effects predicted *in silico*.

Here, we establish the molecular basis of sequence-dependent N-glycosylation by the human OST, revealing how individual residues within the acceptor sequon control substrate recognition and glycosylation efficiency in human cells. By integrating MD simulations and alchemical free-energy calculations with cellular glycosylation assays, we link sequon composition to binding energetics and productive catalytic positioning, providing mechanistic principles for the prediction and engineering of N-glycosylation sites.

## Results

### Global structural assessment

To gain molecular insight into the stability and key interactions of the Michaelis complex (MC), classical MD simulations were performed as described in the Methods section. The initial peptide ligand was modelled with the sequence ^Ace-Nt^-A-Y-A-N-A-T-S-A-A-^Ct-Nme^ and the LLO ligand used in our simulations consists of the well conserved 11-residue oligosaccharide structure, Glc3Man6GlcNAc2, attached to a biphosphate moiety linked to the lipid chain (see Figure S1).

Figure 1. The STT3 subunit is given in the light green cartoon representation; peptide ligand is depicted in magenta and LLO is shown with the light-yellow sticks. The N-terminal region of STT3 forms a transmembrane (TM) domain consisting of 13 TM helices, while the C-terminal region adopts a soluble, globular fold, facing the ER lumen. Consistent with this structural organization, the TM region was submerged in the lipid bilayer generated by Packmol-Memgen and the globular domain was left exposed to the solvent (see Figure 1a). The active site is located in the C-terminal domain (see Figure 1b). This C-terminal domain of the STT3 subunit contains two conserved D*X*D motifs located in the first and second external loops, corresponding to EFD (residues 47-50) (see Figure 1c on the top) and DNE (residues 167-169) (see Figure 1c on the bottom). The C-terminal domain also contains an SVSE motif (residues 348-351) in the fifth external loop (see Figure 1c), which correspond to the conserved TIXE motif found in Archaea and Eubacteria.^33^ All these conserved motifs are summarized in Figure S4. Previous studies on archaeal and bacterial OSTs have shown that conformational changes in the N- and C-terminal halves of this fifth loop are essential for binding of the LLO and the acceptor sequon, respectively.^33,62,63^ As shown in Figure 1a, aliphatic tail of the LLO is accommodated within the hydrophobic surface of the membrane-inserted subunit, while the oligosaccharide moiety extends into the C-terminal and aqueous, hydrophilic regions. As mentioned, our initial model was constructed based on the cryo-EM structure (PDB ID: 6S7O^21^) lacking LLO. The LLO structure was taken from the yeast oligosaccharyltransferase complex (PDB ID: 8AGC^42^) after structural alignment. In addition, the 6S7O structure contains a Mn^2+^ ion, which was replaced by Mg^2+^ in our model. Therefore, to assess the structural accuracy, our model was compared to the later-released cryo-EM structure of a OST complex, containing GRP94 xaperone bound to the secretory translocon in complex with CCDC134 and FKBP11 subunits (PDB ID: 9N9J).^22^ Since no divalent metal ion is resolved in the 9N9J active site, the comparison with our model was focused on the conserved backbone architecture and LLO binding mode. Our model and the LLO binding mode show close agreement with the new experimental structure (see Figure S5). The RMSD over all common atom-pairs is 0.72 Å, and 0.24 Å within 10 Å of the anomeric carbon, consistent with a well-converged local structure. A slight displacement observed in the biphosphate and sugar units (Figure S5, right panel), likely reflects limited local resolution and the shorter glycan in the experimental structure (two sugar units versus the more extended glycan in our model).

**Figure 1:**
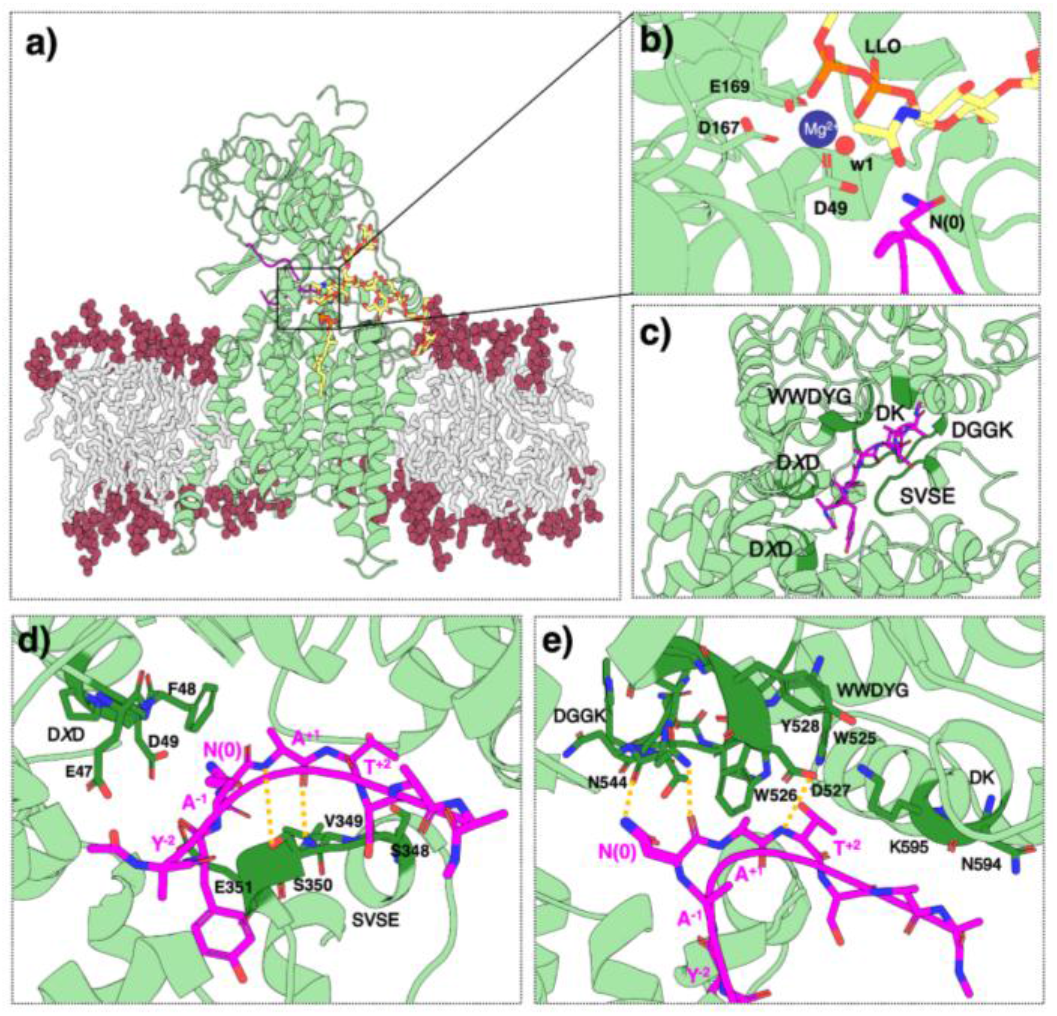
Structural model of the STT3-LLO-peptide complex embedded in the membrane. **(A)** Cross-section of the STT3 protein shown in cartoon representation (light green) embedded in a POPC lipid bilayer (grey sticks with red spheres for the phosphate headgroups). The peptide ligand is depicted in magenta, with the nucleophilic Asn residue N(0) shown in stick representation. The lipid-linked oligosaccharide (LLO) is shown with the light-yellow sticks. **(B)** Close-up view of the active site highlighting Mg^2+^ coordination. The Mg^2+^ ion (blue sphere) is bidentately coordinated by the LLO biphosphate moiety and coordinated by D49, D167, E169 and one water molecule. **(C)** Overall view of all STT3 motifs (darker green color) engaged in the interaction with the peptide ligand given in magenta. **(D)** Close-up of N-terminal domain motifs, including first D*X*D and SVSE, showing their interactions with the ligand. **(E)** Close-up of C-terminal domain motifs (DGGK, WWDYG and DK) and their interactions with the peptide ligand. Key residues within each motif are shown in stick representation. In panels d) and e) orange dashed lines indicate hydrogen-bond interactions discussed in the main text.

In a recently published study on the yeast OST, positional restraints were applied during the entire production time on the acceptor Asn side chain, position of the metal ion, protein backbone and of the LLO heavy atoms in order to preserve the integrity of the catalytic architecture within a reduced OST model.^34^ While this strategy may improve simulation stability, it can also bias the orientation and local conformational ensemble of the sequon, potentially influencing the inferred glycosylation-competent poses. In contrast, our approach allows the sequon to evolve without such restraints, enabling a less biased characterization of the conformational landscape. Notably, our simulations remained structurally stable throughout the trajectory replicates despite the absence of any positional restraints, supporting the robustness of the observed conformational ensembles. The root mean square deviation (RMSD) and root mean square fluctuation (RMSF) analyses of the three replicas are provided in the Supplementary Information (see Figure S6a-c). In general, STT3 protein shows relatively low flexibility (see Fig S6a). The most flexible regions are located quite far from the sequon pocket (see Figure S6b), including residues 1–14, 284–319, 420–462, 493–498 and 696–705 (yellow bars on the Figure S6a). These flexible regions correspond either to N- or C-terminal segments (1-14 and 696-705) or to the loop regions connecting conserved structural elements (284–319, 420–462 and 493–498). Notably, these segments show missing electron density in the cryo-EM structure (PDB ID: 6S7O^21^), consistent with their intrinsic flexibility. However, given that these regions are distal to the catalytic centre (see Figure S6b), they are not expected to significantly influence the geometry of the active site or the stability of the MC.

### Active site interactions and Mg^2+^ coordination

The donor sugar moiety forms stabilizing interactions with STT3 subunit residues via its hydroxyl groups, while the biphosphate group of LLO remains stably positioned in the active site close to the Mg^2+^ ion. Rather than a water-mediated coordination, as previously suggested in other OSTs ^33^, our model suggests that the biphosphate moiety could coordinate the Mg^2+^ ion directly in a bidentate manner (see Figure 1b). The results of our MD simulations further support this coordination geometry, as reflected by the narrow distribution of distances between the biphosphate oxygen atoms and Mg^2+^ (see Figure S7a-f). These distances remain stable across all three independent replicas and, additionally, no drift was observed. Such a coordination mode is actually commonly observed in enzymes exhibiting phosphatase activity.^64^ In addition to the biphosphate group, our model suggests that Mg^2+^ is coordinated directly by D49 from the first DXD motif (see Figure S7e) and D167 and E169 from the second DXD motif (see Figures S7c and S7d). The distances between these residues and Mg^2+^ remain stable throughout the simulation and all the replicas, supporting a robust coordination. The remaining coordination site is occupied by a water molecule (w1 in Figure 1 while the corresponding distribution is shown in Figure S7f), resulting in a stable octahedral geometry characteristic of Mg^2+^ in biological systems.^65^

Our MD simulations show that the side chain of the acceptor Asn residue in the sequon, Asn(0), is positioned on the β-face of the donor sugar, opposite the sugar-phosphate bond (Figure 1b). This arrangement agrees with observations from yeast OST (PDB ID: 8AGC^42^). The Asn amino group remains within ∼3.2 Å of the anomeric carbon (Figure S8a) while the N–C1–O angle fluctuates around ∼140° (Figure S8b), both consistent with a productive S^N^2 geometry.^66,67^ In fact, the average value of the N–C1–O angle from our MD simulations is somewhat closer to the ideal 180° geometry than reported average from the previous theoretical studies (showing a maxima under 130°).^67^ Additionally, acceptor Asn forms a persistent hydrogen bond with D49 (Figure S8c) from the first DXD motif, that anchors this residue near the donor sugar in the correct orientation, in contrast to previous studies, where no interactions were reported for between side-chain carboxamide group of the acceptor Asn and OST.^22,33^ Thus, in addition to the hypothetical twisted amide mechanism previously proposed for the activation of the inert amide nitrogen, our model suggests D49 could potentially also act as a base during the catalysis, facilitating deprotonation of the Asn amino group in the proposed S^N^2 mechanism.^64^ Further studies are required to fully elucidate the catalytic pathway.

### Substrate peptide-STT3 interactions

The donor sugar moiety forms stabilizing The interactions between N- and C-terminal motifs of the human STT3 and the peptide substrate containing the acceptor sequon in our model are shown in Figure 1, panels d and e, respectively. The observed interactions closely resemble those previously reported for archaeal OST.^33^ These include contacts with the N-terminal region of STT3: SVSE motif and the first DXD motif (see Figure 1c). In particular, sequon recognition has previously been linked to the SVSE motif through inter-chain hydrogen bonds.^40^ Consistently, we also observe interactions between the SVSE motif and the peptide substrate. Specifically, S350 interacts with the +1 position of the sequon via its side-chain hydroxyl group and backbone NH, forming hydrogen bonds (Figure 1d). Our MD simulations also reveal transient interactions of E351 with the -1 and +1 positions of the substrate peptide, but most importantly with the backbone NH of Asn(0), suggesting a contribution to the stabilization of the nucleophile. Interestingly, a recent study on yeast OST suggested that its E350 residue acts as a conformational switch that senses sequon complementarity and may serve as an assisting base during catalysis.^34^

On the other side, C-terminal interactions of the STT3 subunit with the sequon involve the DGGK (also known as DN*X*T*Z*N*X*), WWDYG, and DK motifs, which have also been recognized as important in archaeal OSTs.^33^ In the human OST, the DGGK motif corresponds to the sequence DNNTWNN (residues 543–549, see Figure S4). The side chain of N544 from this motif forms a stable hydrogen bond with backbone of the acceptor Asn of the sequon (distance 1.9 ± 0.2 Å), keeping the nucleophile well-positioned (see Figure 1e). The backbone carbonyl oxygen of N544 also forms a hydrogen bond interaction with the carboxamide group of the nucleophile (distance 2.5 ± 0.4 Å), suggesting a key role in nucleophile orientation (see Figure 1e). The WWDYG motif, spanned by residues 525–529, also appears to be of a great importance for sequon recognition (see Figure 1e). In particular, W526 interacts with the +1 position (Ala) via CH–π interactions, while side chain of the D527 forms strong contacts with both the backbone NH and the hydroxyl side chain of the +2 residue of the sequon (Thr). Finally, the DK motif (Figure 1e), in this case represented by NK (residues 594-595), positions K595 in very close proximity to the peptide, contributing to substrate stabilization.

### Relative binding free energies of mutants

Accurate prediction of N-glycosylation efficiency by sequence-based models depends critically on optimizing the glycosylation acceptor sequence, in particular its neighbouring residues. In the present study, we explore this effect through changes in binding free energies and their correlation with the experimentally measured glycosylation efficiencies from our *in cellulo* experiments. Specifically, to quantitatively assess the effects of single-point mutations on glycosylation efficiency, we hypothesized they would be reflected in changes in the binding free energies of the sequons. Higher binding affinities should correlate with higher N-glycosylation rates. To determine the relative binding free energies associated with each mutation, alchemical free-energy calculations were performed as described in the Methods section. As the reference acceptor polypeptide to determine free energy changes we used the nonapeptide sequence ^Ace-Nt^-A-Y-A-N-A-T-S-A-A-^Ct-Nme^, with the central Asn residue defined as position 0 (fourth residue in the peptide). Positions 0, +1 and +2 were defined relative to this acceptor glycosylation sequon. In parallel to the free energy changes, 500 ns of classical MD simulations were conducted for each mutant complex to provide a structural framework for interpreting the trends observed in the computed free energies. Averaged free energy changes for each mutation are listed in Table 1. Detailed free energy changes for each alchemical transformation in aqueous and protein environment and within each replica are given in Table S1. Some key interactions (de)stabilizing each mutant within the OST-A active site are shown in the corresponding panels of Figure 2.

**Table 1:** Relative binding free energies (ΔΔG, kcal·mol^−1^) associated with each mutation. Free energy changes are calculated as differences between the free energy change in water and protein environments, averaged over five independent replicas. Detailed values for each alchemical transformation in both environments and all five replicas are provided in Table S1.

| Variant | $\Delta G_{\text{water}}$<br>(kcal·mol<sup>-1</sup>) | $\Delta G_{\text{protein}}$<br>(kcal·mol<sup>-1</sup>) | $\Delta\Delta G_{\text{total}}$<br>(kcal·mol<sup>-1</sup>) |
| --- | --- | --- | --- |
| N<sub>0</sub>Q<br>(QAT) | $0.68 \pm 0.07$ | $3.53 \pm 0.22$ | $2.85 \pm 0.24$ |
| A<sub>+1</sub>S<br>(NST) | $-10.21 \pm 0.01$ | $-9.80 \pm 0.13$ | $0.41 \pm 0.14$ |
| A<sub>+1</sub>F<br>(NFT) | $-4.86 \pm 0.02$ | $-4.74 \pm 0.15$ | $0.12 \pm 0.16$ |
| A<sub>+1</sub>L<br>(NLT) | $-0.81 \pm 0.03$ | $-0.04 \pm 0.33$ | $0.77 \pm 0.34$ |
| A<sub>+1</sub>W<br>(NWT) | $-10.43 \pm 0.04$ | $-10.46 \pm 0.54$ | $0.03 \pm 0.55$ |
| A<sub>+1</sub>P<br>(NPT) | $19.31 \pm 0.27$ | $-21.73 \pm 0.25$ | $2.42 \pm 0.37$ |
| T<sub>+2</sub>S<br>(NWS) | $1.13 \pm 0.14$ | $3.56 \pm 0.40$ | $2.43 \pm 0.43$ |
| T<sub>+2</sub>C<br>(NWC) | $8.48 \pm 0.04$ | $12.11 \pm 0.18$ | $3.63 \pm 0.19$ |

**Figure 2:**
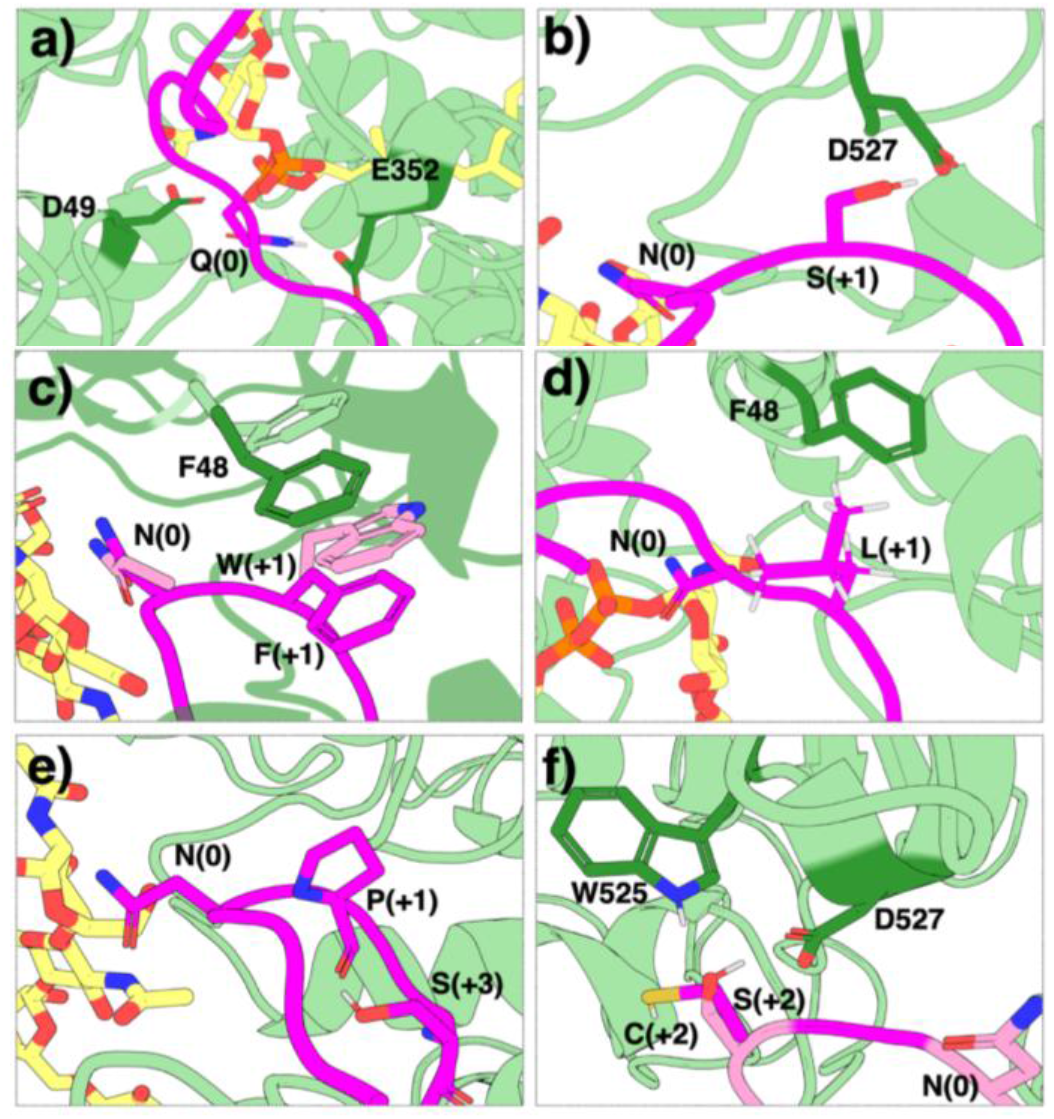
Representative structures showing interactions of the different acceptor sequons within the active site of the human oligosaccharyltransferase OST-A subunit. OST-A is shown in light green cartoon representation and substrate peptide is depicted in magenta, with the important residue depicted in the stick representation. Panels correspond to the following sequon structures: **(A)** N_0_Q; **(B)** A_+1_S; **(C)** A_+1_F and A_+1_W; **(D)** A_+1_L; **(E)** A_+1_P; **(F)** T_+2_S and T_+2_C.

To experimentally assess the influence of the sequons on N-glycosylation efficiency in its native biological environment, a glycosylation mapping assay based on the *Escherichia coli* leader peptidase (Lep) system was envisioned in human cell cultures.^8,38,39^ This well-established approach enables qualitative and semi-quantitative assessment of the OST-driven protein N-glycosylation efficiency from the MD explored candidate sequons. The Lep construct consists of an extended N-terminal luminal domain, two transmembrane helices (H1 and H2) separated by a cytoplasmic loop (P1), and a large C-terminal (P2) luminal region. Two N-linked glycosylation acceptor sites (G1 and G2) were introduced into the N- and C-terminal luminal domains, respectively.^8,38,39^

Because N-linked glycosylation occurs exclusively within the lumen of the endoplasmic reticulum (ER), where the active site of the OST is located, the glycosylation pattern provides a reliable readout of protein N-glycosylation. Each glycosylation event increases the apparent molecular weight of the protein by approximately 2.5 kDa, allowing mono- and doubly-glycosylated species to be readily distinguished by gel electrophoresis and Western blot analysis when adding a proper tag into the Lep vehicle.

In our experimental setup, the first glycosylation acceptor site (G1) was maintained as a canonical NST sequon and served as an internal control for efficient ER targeting and membrane insertion of the Lep reporter construct. Therefore, G1 provides a reliable indicator that the reporter protein reaches the ER lumen and adopts a topology compatible with OST-mediated modification. Mutations were introduced exclusively at the second glycosylation acceptor site (G2), allowing the glycosylation efficiency of this site to be assessed independently of differences in ER targeting or membrane insertion. Consequently, the glycosylation pattern observed at G2 directly reflects the effect of the introduced mutations on the tested sequon recognition and modification by the OST (Figure 3a-c).

**Figure 3:**
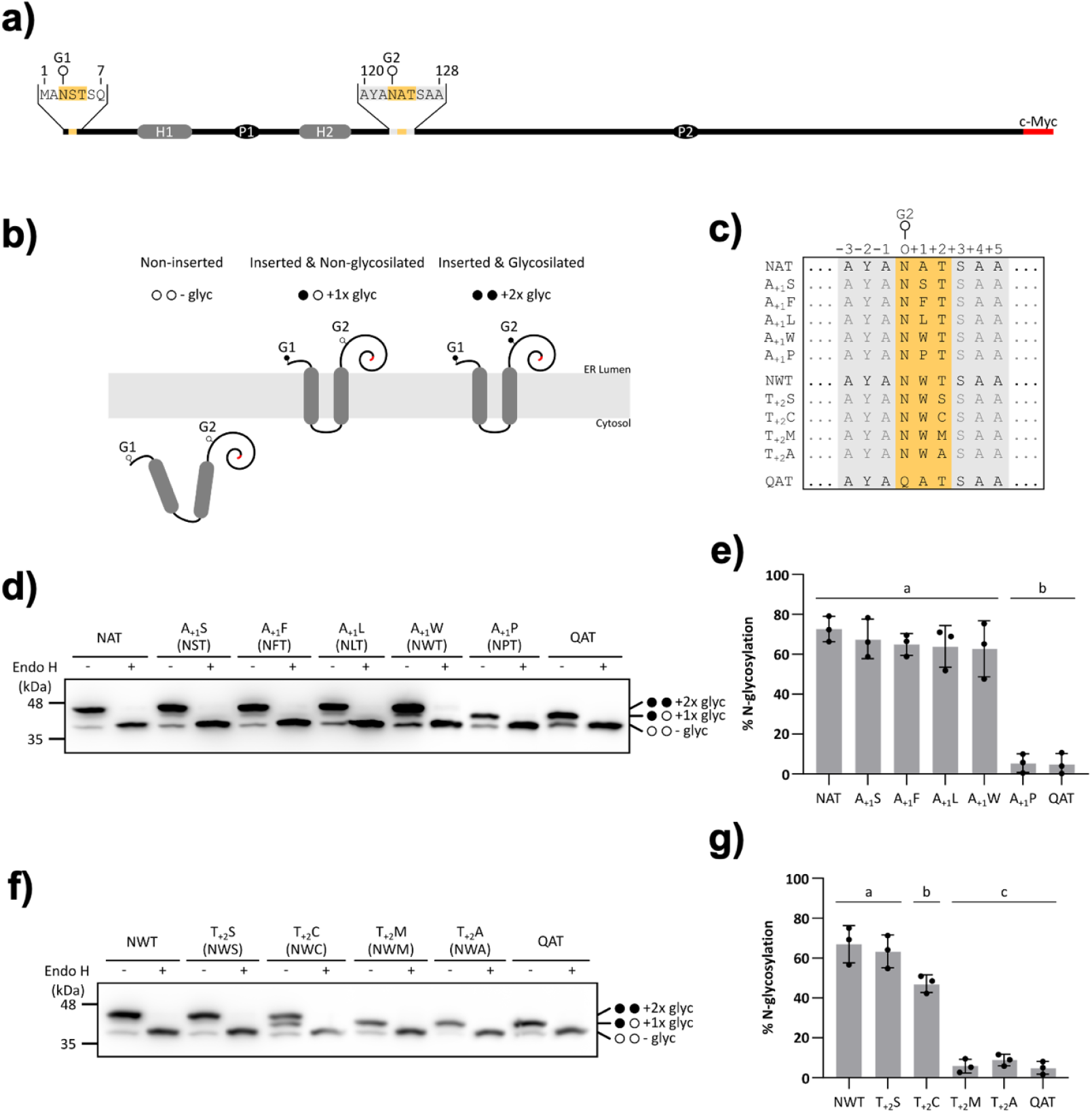
N-glycosylation efficiency of Lep mutants with different N-glycosylation sequons in HEK 293T cells. **(A)** Schematic of the *Escherichia coli* leader peptidase (Lep) construct used as a reporter protein to explore N-glycosylation. The construct contains two transmembrane segments (H1 and H2, highlighted in grey), an unmodified N-terminal glycosylation site (G1), a second acceptor site (G2) where different N-glycosylation sequons were tested and a C-terminal c-Myc tag (highlighted in red). **(B)** Summary of the possible N-glycosylation patterns of Lep in the experimental setup. **(C)** Lep mutants tested, where the flanking residues and the tested sequons are highlighted in grey and yellow, respectively. **(D-G)** Western blot analysis of N-glycosylation efficiency of Lep mutants in +1 (d,e) and +2 (f,g) positions in HEK 293T cell lysates, which were treated with Endo-glycosidase H and incubated with anti-c-myc antibody. For each sample, Endo H treated (+) and mock treated (-) were analyzed. Efficiencies were calculated as described in Materials and Methods section and shown as mean percentage and standard deviation of N-glycosylation quantified in the Western blot analyses from independent triplicates for +1 (e) and +2 (g) positions. Groups labeled with letters a, b and c are significantly different according to one-way ANOVA followed by Tukey’s test for multiple comparisons (p < 0.0001).

To specifically investigate and validate the impact of the G2 acceptor sequons on N-glycosylation efficiency, the initial Lep sequence containing the second glycosylation site (^120^P-T-L-N-S-T-D-F-I^128^) was replaced with the engineered peptide sequence used in our MD simulations (^120^A-Y-A-N-A-T-S-A-A^128^). Thus, the original G2-containing region was substituted by the designed sequence while preserving the overall Lep reporter framework. This G2 site was positioned well away from the C-terminus, (separated by 238 amino acids) thereby preventing potential bypass of glycosylation by the OST.^18,25^ Using this experimental system in human HEK 293T cells, the N-glycosylation efficiencies of each sequon variant were evaluated. To ensure that the increase in molecular weight was due to glycosylation of the acceptor sites, samples were incubated in the presence (+) or absence (−) of Endoglycosydase H (EndoH), a glycan-removing enzyme.

### Position 0

At position 0, Asn is the residue that undergoes glycosylation and is therefore strictly conserved in canonical N-glycosylation sequons. Given the chemical similarity between Asn and Gln, we investigated why glutamine could not support glycosylation in this context. Upon N^0^Q mutation our experimental results show a complete loss of glycosylation efficiency in this QAT sequon (see Figure 3d-g). Consistently, the predicted change in binding free energy is positive, around 2.8 kcal·mol^−1^ (see Table 1), indicating a decrease in binding affinity when Asn is substituted by Gln. However, this free-energy penalty is relatively modest, suggesting that binding effects alone do not explain the observed functional loss.

To gain further insight into the loss of activity, we analyzed MD simulations of the QAT variant (see Figure 2a). In the mutant, the longer side chain of glutamine disrupts the anchoring interaction with D49 of the DXD motif (Figure S9a) and reorients toward E351 of the conserved SVSE motif (Figure S9b). Although the interaction with E351 partially restores stabilization of the side-chain amide, this alternative arrangement does not preserve its interaction with N544 of the DGGK motif observed for the canonical Asn (see Figure 1e). As a consequence of this reorganization, the nucleophilic attack distance between the anomeric carbon of the sugar and the side-chain nitrogen atom of the acceptor residue shifts from a catalytically competent distribution centered at ∼3.2 Å in the canonical Asn to ∼6.7 Å in the N^0^Q mutant (see Figure S9c), placing the system outside the reactive configuration in a pose incompatible with efficient catalysis.

### Position +1

To assess the contribution of the +1 position to N-glycosylation efficiency, single-point mutations were introduced at the position immediately following the acceptor Asn. Alanine, used as the reference, was mutated to S, F, L, W and P (A_+1_S, A_+1_F, A_+1_L, A_+1_W and A_+1_P). The relative binding free energies are shown in the Table 1.

Except for A_+1_P, both our calculations and *in cellulo* experiments support a broadly permissive role for the +1 position for hydrophobic and polar residues. The calculated relative binding free energies show no significant preference among the A_+1_S, A_+1_F, A_+1_L, and A_+1_W variants, with all values remaining within or close to the estimated error bars (Table 1). Thus, the free-energy simulations indicate that aromatic residues (F and W), a bulky hydrophobic residue (L), and a polar residue (S) are all well tolerated at this position. This conclusion is in good agreement with our *in cellulo* assays, which also show minimal discrimination among these mutations (Figure 3d,e and S10a). Therefore, in contrast to previous studies reporting pronounced sequence preferences at the +1 position,^24^ our combined computational and cellular results suggest that this position is relatively permissive, provided that proline is excluded.

The lack of tolerance for Pro at the +1 position, captured in our *in cellulo* experiments, is likely related to structural features previously described for archaeal OSTs.^33^ Proline imposes strong conformational constraints on the peptide backbone. In our MD simulations, the backbone ϕ dihedral angle of Pro was measured to be around –70°, whereas Ala, for example, sampled substantially lower ϕ angles (between -140° and -170°) (see Figure S11a). Similar finding has previously observed in the structural study of the eubacteria and euryarchaeon OSTs, where residues within the sequon preferentially adopt ϕ values substantially lower than -70°.^33^ In addition, in comparison to alanine, proline also lacks the backbone amide hydrogen that can participate in hydrogen bonding in the active site stabilizing the enzyme-peptide complex. Notably, this destabilization is partially compensated by a stabilizing intra-peptide interaction with the +3 Ser side chain (see Figures 2e and S11b). As a result of this balance between lost and compensating interactions, the A_+1_P mutation showed a moderate relative free energy penalty (∼2.4 kcal·mol^−1^, see Table 1). Although this value indicates reduced affinity, it does not explain why glycosylation is completely abolished experimentally. This apparent discrepancy suggests that binding free-energy differences alone do not fully capture the structural consequences of proline incorporation at the +1 position. Indeed, the reorganized interaction network promotes a narrower and substantially altered sequon geometry. This is reflected in the RMSF profile of the original and mutated sequons (see Figure S11c and in the backbone dihedral angle distributions (see Figure S11a). At position +2, the φ and ψ distributions no longer overlap with those of the wild type, indicating a substantial reorganization of the accessible backbone conformations. More importantly, this conformational rearrangement leads to an unproductive catalytic pose: in the WT ensemble, the Asn–anomeric carbon distance samples near-attack conformations (<3.5 Å) in ∼70% of frames, whereas the A_+1_P mutation reduces this population to ∼5% (see Figure S11d) and at the same time the N– C1–O angle distribution is also perturbed from a near-ideal ∼140° toward a distorted ∼90° alignment (see Figure S11e). Collectively, these findings indicate that the A_+1_P mutation abolishes glycosylation not simply by destabilizing substrate binding, but primarily by trapping the sequon in a catalytically unproductive conformational state.

Although bulky and aromatic residues, such as phenylalanine (A_+1_F) and tryptophan (A_+1_W), at the +1 position may initially appear unfavourable for glycosylation, our experiments and free energy calculations suggest that both are well tolerated (see Figures 2c, 3d,e and Table 1). Specifically, our MD simulations of the mutants explain this observation, suggesting that their aromatic side chains can be stabilized by π–π stacking interactions with the nearby F48 residue of the first DXD motif in STT3. This likely offset the steric penalty associated with their bulky nature, resulting in very minor free energy penalties for binding (0.12 ± 0.16 and 0.03 ± 0.55 kcal·mol^-1^, respectively).

In the case of aliphatic residues, our relative binding free energy calculations suggest that the hydrophobic leucine also remains energetically viable at the +1 position, with a free-energy penalty of only 0.77 ± 0.34 kcal·mol^-1^, as confirmed also in our experimental glycosylation rates. Our MD simulations indicate that Leu can easily be accommodated by a local hydrophobic pocket but also by a CH–π interactions with F48 of the DXD motif (see Figure 2d), thereby enabling stable packing interactions without introducing steric clashes or disrupting the local interaction network. Similarly, our relative binding free energy calculations suggest that a polar serine is also well accommodated at the +1 position, the free energy difference being even smaller (0.41 ± 0.14 kcal·mol^-1^, see Table 1) as also corroborated in the glycosylation experiments (Figure 3d,e). Analysis of the simulation trajectories indicates that the serine hydroxyl group engages in dynamic, hydrogen-bonding interactions with D527 of the WWDYG motif (Figure 2b), contributing to local stabilization of the acceptor peptide. Additionally, the side chain of serine also remains partially exposed to solvent, enabling additional stabilization through transient interactions with bulk water molecules accessing the opposite side of the sequon.

Taken together, our experimental and computational results suggest that the +1 position of the sequon is relatively permissive toward chemically diverse residues, as among the variants investigated here and only proline showed a clear unfavourable effect on the N-glycosylation rate.

### Position +2

The amino acid at the +2 position of the N-glycosylation sequon (Asn-X-Ser/Thr) plays a decisive role in substrate recognition by the OST complex, yet the structural and chemical constraints governing this specificity remain incompletely understood. Previous *in vitro* studies have proposed that threonine-containing sequons serve as superior glycosylation substrates relative to serine, due to a stereo-electronic advantage and the participation of the hydroxyl group in a proton relay that facilitates transfer of the oligosaccharide to the Asn β-amide.^24,29^ Inspired by recent evidence that even non-canonical motifs such as Asn-*X*-Cys can occasionally be glycosylated in mammalian cells,^30,31,68^ we systematically explored how the nature of the residue at +2 affects glycosylation efficiency and fidelity. To this end, we designed a panel of five sequon variants: NWT, NWS, NWC, and two sulphur containing and non-polar controls: NWM and NWA. We fixed the +1 position as Trp based on the findings of Kasturi et al. (1997),^24^ who demonstrated that Ser-to-Thr substitutions at the +2 position produced some of the largest changes in N-glycosylation efficiency when Trp was present at +1. This enabled us to isolate the contribution of the +2 residue while keeping the +1 constant.

We examined the glycosylation behaviour of these sequons in the human cells and by computational free energy calculations. In contrast with the position +1, our *in cellulo* assays showed a clear hierarchy in glycosylation efficiency depending on the residue at +2 position: NWT > NWS ≫ NWC, while NWM and NWA remained unglycosylated (see Figure 3f,g and S10b). The calculated relative binding free energies (see Table 1) show exactly the same trend, with T_+2_S exhibiting a moderate penalty (∼2.4 kcal·mol^−1^) and T_+2_C a larger destabilization (∼3.6 kcal·mol^−1^), consistent with the experimentally observed glycosylation efficiency for the 3 sequons (NWT > NWS ≫ NWC). Analysis of the structural models show that in the NWT sequon both the main chain amino and side chain hydroxyl groups of Thr are donating hydrogen bonds to the D527 residue of the WWDYG motif (Figures 2f and S12b-c). Simultaneously, the oxygen atom of the hydroxyl group accepts a hydrogen bond from W525 of the same WWDYG motif (see Figure S12d). The mutation of Thr to Ser removes the β-methyl group, broadening and shifting the torsional distribution of the hydroxyl side chain (reflected in the values of the χ dihedral, see Figure S12a). This increased conformational flexibility reduces hydrogen-bond directionality and weakens interactions persistence. This effect is further amplified when Thr is mutated to Cys. The T_+2_C mutant does not only lacks the β-methyl group, but also replaces the hydroxyl with a thiol, a weaker hydrogen-bond donor due to the lower electronegativity of sulphur compared to oxygen. This is also reflected in the χ angle distribution of the thiol side chain, which is markedly broader and shifted with respect to Thr and Ser (see Figure S12a). These structural changes provide a rationalization for the observed trends in the glycosylation selectivity towards position +2.

### Human N-glycosylation site analysis

Next, we generated a human N-glycosylation site dataset by selecting experimentally validated glycosites from the N-GlycositeAtlas database^16^ that matched the canonical N-X-S/T (X ≠ P) sequon and overlapped with reviewed human N-glycosylation site annotations in UniProt. This dataset was used to investigate amino acid preferences at positions flanking the glycosylated asparagine, assess the prevalence of variants of the canonical N-X-S/T sequon and examine the influence of the distance between the glycosylated asparagine and both the protein N- and C-termini.

We first analysed amino acid enrichment at positions surrounding the glycosylated asparagine to identify sequence preferences beyond the canonical N-X-S/T sequon. Inspection of the complete amino acid enrichment profile across a 41-residue window centred on the glycosylated asparagine (20 residues on either side) revealed that positions −2, −1, +1, and +3 exhibited the largest deviations from the overall amino acid frequencies (Figure S13a). These four positions have also been the focus of most previous experimental studies (Table S2) and are included within the nonapeptide analysed in our computational and experimental work; accordingly, they are shown in detail in Figure 4a.

**Figure 4:**
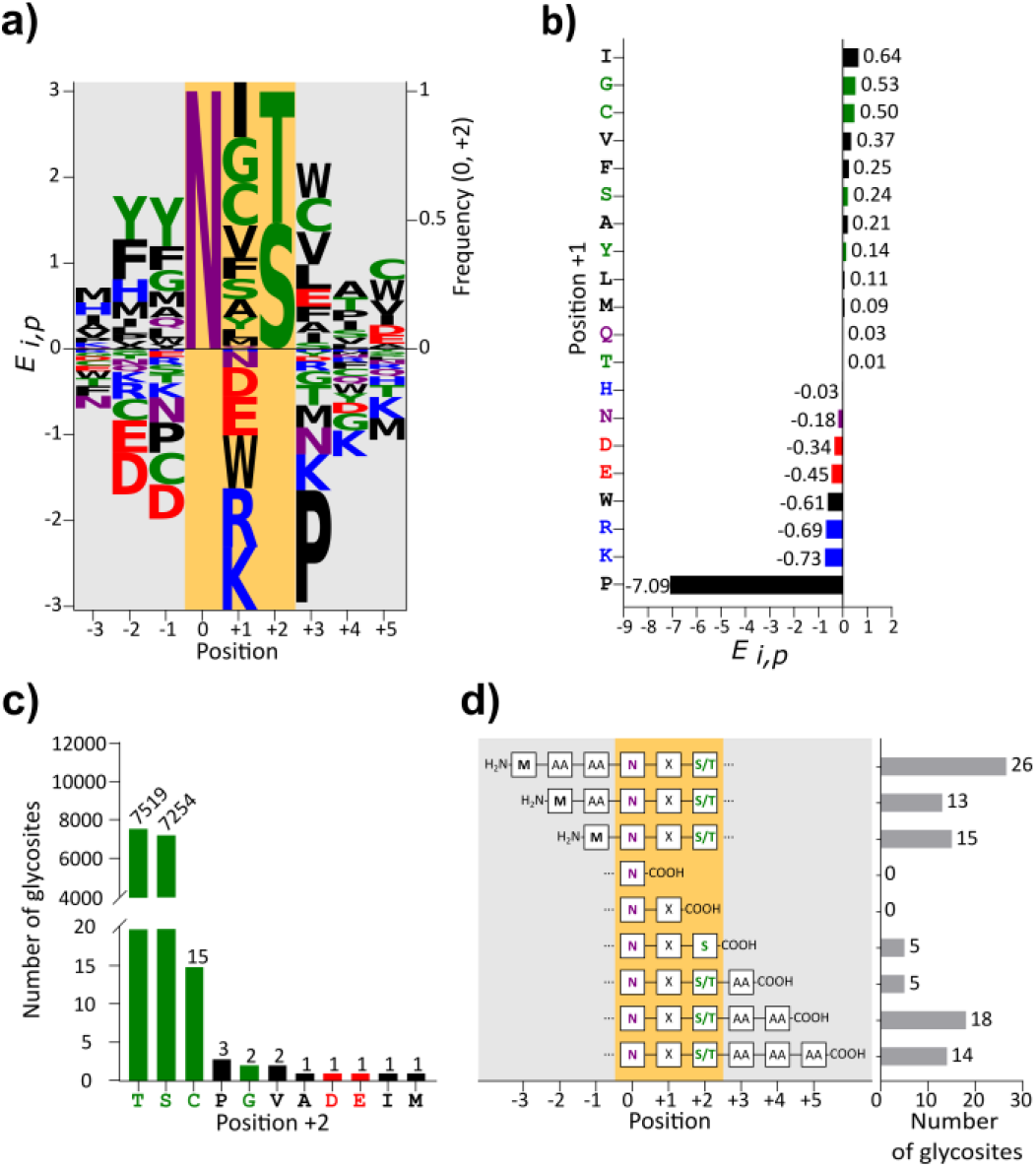
Sequence analysis of the 14.800 glycosites dataset. **(A)** Sequence logo showing amino acid enrichment of -3, -2, -1, +1 (proline not shown), +3, +4, and +5 positions and amino acid frequency of 0 and +2 positions. Positions 0, and +2 were omitted from the enrichment analysis because of the predominance of asparagine residue at the 0 position, and serine and threonine residues at the +2 position. **(B)** Amino acid enrichment at the +1 position. **(C)** Amino acid distribution at the +2 position. **(D)** Distribution of glycosites according to their proximity to the N- and C-termini of the protein. Amino acid residues are colored according to WebLogo3 chemistry color scheme. Polar-green (G,S,T,Y,C); neutral-purple (Q,N); basic-blue (K,R,H); acidic-red (D, E); and hydro-phobic-black (A,V,L,I,P,W,F,M).

As expected, position +1 displayed the strongest enrichment pattern. Polar and hydrophobic residues (I, G, C, V, F, S, A, Y, L and M) were positively enriched, whereas charged residues (D, E, R and K), together with tryptophan, were markedly underrepresented (Figure 4A). This broad tolerance toward chemically diverse residues is consistent with our computational and *in cellulo* results, which show that the +1 position can accommodate polar, aliphatic and aromatic side chains through alternative stabilizing interactions within the STT3 binding pocket.Notably, only six of the 14,800 glycosites contained proline at position +1, corresponding to an enrichment score of −7.09 (Figure 4B), in agreement with the complete loss of glycosylation observed for the A+1P variant in our cellular assays and with the structural disruption revealed by our MD simulations. The enrichment pattern observed at this position recapitulates previously identified determinants of glycosylation efficiency (Table S3). Specifically, experimental studies have shown that glycosylation efficiency is reduced when tryptophan or proline occupies the +1 position^11,24,27^ whereas Kasturi et al. and Shakin-Eshleman et al. further demonstrated that aspartate or glutamate at this position also decrease glycosylation efficiency. The apparent discrepancy for tryptophan, which is well tolerated in our model sequon but underrepresented in the human glycosite dataset, further suggests that preferences at the +1 position are strongly modulated by the surrounding sequence context.

At the +2 position, 99.82% of glycosites contained either serine or threonine, whereas only 27 carried a different amino acid. Among these atypical sequons, 15 contained cysteine (Figure 4c). Our experimental and computational analyses, together with previous studies, indicate that cysteine can support N-glycosylation in specific sequence and structural contexts, giving rise to non-canonical NXC glycosylation motifs. However, these events appear to be exceptional compared with the canonical NXT and NXS sequons. Both our molecular dynamics simulations and experimental validation indicate that NXT sequons provide the most favourable atomic-level environment for glycosylation, followed by NXS, whereas NXC motifs provide a substantially less favourable environment for efficient glycan transfer. The structural changes discussed above and the redox reactivity of the cysteine thiol group are likely to reduce the accessibility and/or stability of NXC motifs *in cellulo*.

Together, these findings provide a mechanistic explanation for the strong evolutionary preference for threonine and serine over cysteine at the +2 position of N-glycosylation sequons.

Positions −2, −1, and +3, which flank the canonical sequon, also exhibited pronounced amino acid preferences. Positions −2 and −1 were enriched in aromatic residues, particularly tyrosine and phenylalanine, indicating that the sequence preferences observed across the human glycoproteome recapitulate previously identified determinants of glycosylation efficiency (Table S2). Specifically, experimental studies have shown that phenylalanine at position −1^26^ and both tyrosine and phenylalanine at position −2^27,28^ enhance glycosylation efficiency. At position −1, aspartate, cysteine, proline and asparagine were underrepresented, in agreement with previous reports showing that proline and aspartate reduce glycosylation efficiency.^26,27^ Likewise, negatively charged residues (D and E) were depleted at position −2. At position +3, proline was again strongly depleted, consistent with the marked reduction in glycosylation efficiency reported by Mellquist et al., whereas tryptophan, cysteine, glutamate, and the hydrophobic residues valine and leucine were enriched. Finally, analysis of asparagine frequencies revealed that this residue was most strongly depleted at positions −1, +1, and +3 (Figure S13b), whereas no asparagine residues were observed at position +2 (Figure 4c), presumably to prevent the formation of adjacent glycosites.

## Discussion

In this work, we combined atomistic molecular dynamics simulations, alchemical free-energy calculations and *in cellulo* glycosylation assays to investigate the molecular determinants governing sequence-dependent N-glycosylation by the human OST complex. We first generated an atomistic model of the catalytic STT3 subunit in complex with both the lipid-linked oligosaccharide donor and an acceptor peptide substrate starting from the cryo-EM structure complemented with AlphaFold modelling. Our simulations revealed a stable active-site architecture involving direct bidentate coordination of the Mg^2+^ ion by the LLO biphosphate group and conserved residues of the nearby structural motifs. Importantly, the acceptor peptide remains stably accommodated in the catalytic pocket, with the acceptor Asn side chain oriented toward the anomeric carbon of the donor sugar and supported by a network of interactions with conserved STT3 motifs. This binding mode provides a structural framework to rationalize how local sequon variations modulate productive substrate positioning and, ultimately, glycosylation efficiency. To our knowledge, this represents the first well-resolved atomistic model of the human OST with both donor and acceptor substrates simultaneously accommodated in a catalytically competent state, establishing a molecular framework for dissecting substrate recognition and catalysis in the human enzyme.

To assess the contribution of sequon residues to N-glycosylation efficiency, we experimentally tested mutations at positions 0, +1 and +2 using a Lep-based reporter assay in human cells, and computed their effects on binding to STT3 through relative binding free-energy calculations. At position 0, substitution of the acceptor Asn by Gln abolished glycosylation experimentally, while resulting in a moderate binding free-energy penalty, suggesting that the main effect of this mutation is likely related to disruption of the catalytic geometry rather than to loss of binding alone. At the +1 position, both the experimental data and the free-energy calculations indicate a broadly permissive behaviour toward chemically diverse residues, with the clear exception of proline. This permissiveness likely reflects the versatility of the local STT3 binding environment, where residues from the DXD, SVSE and WWDYG motifs provide alternative stabilizing interactions for different side-chain chemistries, including aromatic contacts with bulky residues, hydrophobic or CH–π interactions with apolar residues, and transient hydrogen bonds with polar residues. In contrast, mutations at the +2 position revealed a well-defined hierarchy in glycosylation efficiency, Thr> Ser >> Cys, consistent with the lower electronegativity of sulphur compared to oxygen, with the calculated destabilization of binding and with the importance of maintaining productive interactions with the conserved WWDYG motif of STT3.

Molecular dynamics simulations provided a structural interpretation of these effects, showing how individual mutations can be accommodated by substrate repositioning, changes in the hydrogen-bonding patterns and in the conformational ensemble of the bound sequon. Our results indicate that glycosylation efficiency is governed in terms of binding affinity and by the ability to preserve catalytically competent conformations within the active site. Thus, the simulations connect local sequon composition with substrate recognition, productive positioning and, ultimately, glycosylation efficiency.

A major strength of our experimental approach is that glycosylation efficiency is measured directly in human cells, preserving the complex cellular environment in which the OST machinery operates. This provides a biologically relevant complement to *in vitro* microsomal assays, which remain among the most sensitive and quantitatively precise approaches for measuring intrinsic OST-mediated glycosylation efficiencies. At the same time, our *in cellulo* assays are primarily sensitive to relatively large effects, likely reflecting the robustness of the cellular glycosylation machinery. In living cells, glycosylation efficiency emerges from the interplay of multiple factors beyond the intrinsic catalytic preferences of the OST active site, including protein folding and quality-control pathways, endoplasmic reticulum homeostasis and stress responses, differential contributions of STT3A- and STT3B-containing OST complexes, substrate accessibility and other cellular determinants.Moreover, as very recently suggested,^34^ the sequence context surrounding the N-X-S/T sequon motif can substantially modulate glycosylation efficiency. In the present work, we deliberately employed a simplified model sequon flanked predominantly by alanine residues to isolate the specific contributions of positions +1 and +2. While this strategy facilitates mechanistic interpretation, alternative sequence contexts may establish additional intra-peptide contacts, long-range interactions or conformational preferences that further influence substrate recognition and glycosylation outcomes.

Importantly, despite these limitations, the agreement between our molecular dynamics simulations and cellular validations supports the notion that the intrinsic structural properties of the sequon constitute a major determinant of glycosylation efficiency. Our results provide an atomistic explanation for the experimentally observed hierarchy and reveal previously uncharacterized molecular features of the human OST active site, including the coordination environment of the catalytic metal ion, the structural role of conserved STT3 motifs and the mechanistic consequences of substitutions at positions 0, +1 and +2. To place these findings in the context of the broader literature, we additionally compiled all published quantitative studies examining sequon-dependent N-glycosylation efficiencies (Table S2). Collectively, these studies reveal a remarkably dependence on the context beyond Asn-X-Thr/Ser sequons. We therefore propose that the present work provides a mechanistic bridge between decades of biochemical observations and an atomic-level understanding of sequon selectivity by the human OST complex.

## Acknowledgements

Authors acknowledge the financial support from grants PID2024-157213NB-I00 (to IT) and PID2023-152568NB-I00 (to IM) funded by MCIN/AEI/10.13039/501100011033/and by “ERDF A way of making Europe”; and CIPROM/2022/062 (to IM) funded by Generalitat Valenciana. JFBA is thankful for the FPU 2024 program (FPU24/01055 grant). DB is funded by grant PRDVA246050BELD from Asociación Española Contra el Cancer (AECC). Authors also thank computational facilities from the Tirant supercomputer of the University of Valencia and MareNostrum Supercomputing Center in Barcelona (projects QH-2024-1-0005, BCV-2024-3-0017 and BCV-2025-1-0005) for computational resources.

## Author contributions

IT and IM conceived and designed the theoretical and experimental parts of the study, respectively. JFBA carried out all experimental work and experimental data analysis under the supervision of IM. DB analysed experimental data and compiled glycoprotein databases. MA prepared all the computational models, performed all computational calculations and did the theoretical analysis. IT, JJRP and IM acquired funding. MA and JFBA drafted the manuscript and all authors contributed to manuscript revision and approved the final version.

## Materials and Methods

### System preparation for Molecular Dynamics Simulations

The eukaryotic OST is a multisubunit membrane protein complex composed of eight distinct subunits, among which the catalytic core is provided by the STT3 protein.^40^ The structure of the human protein subunit STT3 was taken from the cryo-EM structure of the human oligosaccharyltransferase (OST-A) complex (PDB ID: 6S7O^21^). Missing residues of STT3 were modelled with AlphaFold2.^41^ The coordinates of the lipid-linked oligosaccharide (LLO), peptide substrate and the catalytic ion were taken from the yeast oligosaccharyltransferase complex (PDB ID: 8AGC^42^) after superimposing it with our model. The LLO ligand used in our simulations consists of the well conserved 11-residue oligosaccharide structure, Glc3Man6GlcNAc2, attached to a biphosphate moiety linked to the lipid chain (see Figure S1). The lipid portion is of a polyprenyl-type (isoprenoid), composed of four isoprene units (C-20 isoprenoid). Catalytic ion Mn^2+^ present in the cryo-EM structure was then replaced by Mg^2+^, as expected in the catalytically relevant form. The initial peptidic substrate was extended by adding two alanine residues at both the N- and C-termini of the model peptide (^Nt^-Y-A-N-A-T-S-A-^Ct^) from the yeast OST structure (PDB ID: 8AGC^42^) and then capped with Ace and Nme groups, respectively, yielding the peptidic ligand ^Ace-Nt^-A-Y-A-N-A-T-S-A-A-^Ct-Nme^. Protonation states of titratable residues were assigned using PROPKA 3.0^43^ at pH 8.0. The system was embedded in a POPC (1-palmitoyl-2-oleoyl-sn-glycero-3-phosphocholine) lipid bilayer, as in some of the previous studies,^44^ generated with Packmol-Memgen.^45^ Water molecules were added to ensure at least 12 Å between any protein/substrate atom and the edges of the simulation box. Counterions (Na+ and Cl−) were added to neutralize the system and to achieve a physiological ionic strength of 150 mM.

### Molecular Dynamics Simulations

The protein and the peptide substrate were described using the ff14SB force field^46^, lipid bilayer with the lipid21 force field^47^and water with TIP3P potential.^48^ LLO was parametrized following the non-standard residue parameterization procedure using Antechamber in Amber 24.^49^ Atomic charges were calculated using the restrained electrostatic potential (RESP) method at the HF/6-31G^*^ level of theory.^49^ The parameters of the LLO are available on the GitHub repository (see Data availability section).

MD simulations with periodic boundary conditions were run with the Amber24 GPU version of pmemd.^50,51^ The system was minimized in successive stages: initially, water molecules and hydrogen atoms were minimized. Then the membrane was minimized, followed by the substrate and peptidic ligand minimization. Finally, the entire system underwent minimization. After minimization, the system was heated to 310 K under NVT conditions using a Langevin thermostat and friction coefficient of 1 ps^−1^, followed by a relaxation (1 ns) in the NPT ensemble. Production simulations were carried out in NVT ensemble during 1 μs for each of three replicas, initiated with different random velocities, to guarantee enough sampling. A timestep of 2 fs was used, with SHAKE constraints on hydrogen atoms bond lengths.^52^ All long-range electrostatic interactions were described using the particle mesh Ewald method (PME) ^53^ and the cut-off radius was set to 12 Å. All inputs and parameter files for our molecular dynamic simulations are available on GitHub repository (under Data availability section).

### Relative Binding Free Energy Calculations

Relative binding free energies were calculated using thermodynamic integration (TI) along alchemical transformations, as described in our previous work and elsewhere.^54–56^ Each alchemical transformation was defined to gradually mutate a single residue of the peptidic ligand into another, while keeping the rest of the system unchanged. The transformations were performed both in the solvated state (peptidic ligand in bulk water) and in the protein complex to construct the thermodynamic cycle to determine the relative binding free energies (see Figure S2). For each transformation, twelve λ windows were employed following a Gaussian quadrature scheme (the corresponding weights were 0.00922, 0.04794, 0.11547, 0.20683, 0.31659, 0.43778, 0.56222, 0.68341, 0.79317, 0.88453, 0.95206, 0.99078).

All free energy calculations were performed in Amber24 using a dual-topology approach and a previously reported GPU-accelerated protocol.^49,51,52^ All simulations were conducted at 310 K, using Langevin dynamics (collision frequency 1 ps^−1^). Only the side chains atoms of the mutated residue were included in the soft-core mask, although in the case of the proline all atoms were included in the soft-core potential. Default values were taken for all the parameters controlling softness of the potentials. Each system was first equilibrated for 5 ns within each λ window, followed by a 40 ns production run. To improve sampling efficiency, Hamiltonian replica exchange (H-REX) was attempted between adjacent λ windows every 10 ps during the production phase, resulting in the average acceptance ratios between 0.35 and 0.45. Five independent replicas were performed in both water and protein environments to calculate the average values and statistical uncertainties reported in this paper are calculated as the standard error of mean.

### Plasmids and Reagents

The sequence of a modified version with extended N- and C-terminal of full-length Leader peptidase (“Lep”, UniProt ID: P00803) gene from *Escherichia coli* was tagged with c-myc epitope at its Ct (EQKLISEEDL) and inserted into the mammalian vector pHDM-S^58^ (Addgene #164433) by replacing the sequence of SARS-CoV-2 S protein using Restriction-Free Cloning^59^:

- Forward primer:
ACTTTGGCAAAGAATTCCGCGGGCGGCCGCTCTAGAGCCACCATGGCGAAT
- Reverse primer:
GTCGACGGTATCGATAAGCTTGGATCCTTACAGATCCTCTTCTGAGATGAG-TTTTTGTTCATGGATGCCGCCAATGCG Then, the ^Nt^-A-Y-A-N-A-T-S-A-A-^Ct^ peptide DNA sequence was cloned in the Lep P2 domain (residues 120-128), again by Restriction-Free Cloning:
- Forward primer:
TCAGGTTCGATGATGCCGACTCTGGCTTATGCCAACGCCACATCAG
- Reverse primer
TCTTTAATGCCATAAGCAAACTTCTCTACCAGTGCGGCTGATGTGGCGTT-GGCA

Mutants at positions 0, +1 and +2 were generated by PCR-based site-directed mutagenesis with the Pfu Plus! mutagenesis kit (EurX, Gdańsk, Poland) according to the manufacturer’s instructions. All sequences were verified by DNA Sanger sequencing at Macrogen (Seoul, South Korea). The primer sequences for mutagenesis and DNA plasmid sequences generated in this work are provided as Supplementary Information.

### Mammalian Cell Culture and Transfection

HEK 293T cells (ATTC, CRL-3216) were grown in Dulbecco’s modified Eagle’s medium (DMEM) (Gibco) supplemented with 10% fetal-bovine serum (FBS), penicillin-streptomycin (P/S) (100 U/ml) and amphotericin B (0.5 μg/ml) at 37°C, 5% CO_2_. Cells were seeded in 6-well plates (2 × 10^6^ cells per plate) and transfected after 24 hours. For transfection procedure, 2 μg per well of plasmids encoding c-Myc tagged Lep mutants were added to a mixture of 8 μL of polyethylenimine (PEI) MW 25,000 (1 mg/mL) (Alfa Aesar) diluted in 200 μL of Opti-MEM reduced serum medium (Gibco). Transfection mixture was incubated for 15 min at room temperature and then added dropwise to cultured cells. At 24 hours post-transfection, cells were harvested and washed with PBS buffer. After a short centrifugation (350 rcf for 5 min on a table-top centrifuge) cells were lysed by adding 250 uL of radioimmunoprecipitation assay (RIPA) buffer (30 mM Tris–HCl, 150 mM NaCl, 0.5% Nonidet P-40 and 1% SDS) supplemented with cOmplete EDTA-free protease inhibitors (Roche) and sonicated in an ice bath in a bioruptor (Diagenode) during 15 min.

### Glycosylation-Based Western Blot Assays

Equal amounts of protein were submitted to endoglycosidase H (EndoH) treatment (1-hour incubation at 37ºC) in a final volume of 50 μL of Sample Buffer (62.5 mM Tris-HCl, pH 6.8, 25% glycerol, 2% SDS, 3% DTT and 0.01% bromophenol blue) followed by SDS-PAGE of both treated and untreated samples and transferred into a PVDF transfer membrane (ThermoFisher Scientific). Protein glycosylation status was analysed by Western Blot using an anti-c-myc antibody (Sigma), anti-rabbit IgG-peroxidase conjugated (Sigma), and with ECL developing reagent (GE Healthcare). Chemiluminescence was visualized using an Image Quant TMLAS4000 mini–Biomolecular Imager (GE Healthcare). Bands corresponding to different glycosylation levels were then quantified with Image J program for % Glycosylation calculations. To this end, band intensity corresponding to the doubly glycosylated species (2×glyco) was quantified using the ROI Manager tool. This value was then divided by the total intensity of all detected glycosylated species (2×glyco + 1×glyco + 0×glyco) and expressed as a percentage. Reported percentage of the doubly glycosylated protein was determined as the average of three independent experiments. Note that in all Western blot experiments a ∼39 kDa band appeared as a result of truncated Lep protein (see Figure S3). These bands were not included in the glycosylation quantification analyses.

### Generation of the Glycosite Dataset

The glycosite dataset was generated by integrating glycosylation sites from the N-GlycositeAtlas database^16^ with glycosylation sites annotations from UniProt database. Glycosites from both sources were merged to create a unified, non-redundant dataset for subsequent analyses. Uniprot entries were filtered to include only reviewed human *(Homo sapiens)* proteins containing annotated N-linked glycosylation sites on asparagine residues. Each glycosite was defined as a 41-amino acid sequence centered on the glycosylated asparagine residue. The final dataset comprised 14.800 glycosites, of which 1.722 were unique to Uniprot, 8.911 were unique to the N-GlycositeAtlas and 4.167 were common to both databases.

Amino acid enrichment *E*_*i,p*_ was calculated for all positions except 0 and +2 using the following equation:

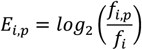

where *f*_*i,p*_ is the frequency of amino acid *i* at position *p*, and *f*_*i*_ is the overall frequency of amino acid *i* across all analysed positions, excluding positions 0 and +2. Positions 0 and +2 were excluded from the enrichment analysis because they are strongly constrained by the canonical N-X-S/T sequon.

### Protein Sequence Alignment

Multiple sequence alignments of the STT3A, STT3, AGLB1 and PGLB protein sequences were generated using Clustal Omega algorithm.^60^ The resulting alignments were visualized using Jalview^61^ and are available in the Supplementary Information (Table S4), while the alignment of the catalytic subunits is given in the Figure S4 of the Supplementary Information. Phylogenetic relationships and pairwise sequence identity were calculated from the Clustal Omega alignment.

