## Supplementary Information for "Molecular Determinants of Sequence-Dependent N-glycosylation by the Human Oligosaccharyltransferase"

### Supplementary Information (SI)

***Index***

**S1.** Chemical and schematic (SNFG) representations of the lipid-linked oligosaccharide (LLO) used in MD simulations (Figure S1) 3

**S2.** Schematic representation of the thermodynamic cycle on the alchemical transformations (Figure S2) 3

**S3.** Full Western blot membranes corresponding to Lep mutants at position +1 and +2 (Figure S3) 4

**S4.** Sequence conservation of the catalytic subunits of oligosaccharyltransferases from representative eukaryotic and prokaryotic organisms (Figure S4) 4

**S5.** Structural alignment of the model with the cryo-EM structure (PDB: 9N9J) of the OST complex (Figure S5) 5

**S6.** Flexibility and stability of the OST-A model during molecular dynamics simulations. (Figure S6) 5

**S7.** Distribution of the distances between the $Mg^{2+}$ and the coordinating residues as depicted in the panel in the middle (Figure S7) 6

**S8.** Distributions of reaction-relevant geometries in the catalytic site during MD simulations (Figure S8) 6

**S9.** Effects of the N0Q mutation on catalytically relevant interactions within the OST active site (Figure S9) 7

**S10.** Quantification of experimental N-glycosylation efficiency for the +1 (a) and +2 (b) positions (Figure S10) 7

**S11.** Effect of the A+1P mutation on the sequon backbone conformation, flexibility and near-attack geometries (Figure S11) 8

**S12.** Geometric determinants relevant for the +2 sequon recognition in the OST-A active site during MD simulations (Figure S12) 9

**S13.** Position-specific amino acid enrichment across the 41-residue glycosite window (Figure S13) 10

**S14.** Free energy changes associated with the alchemical transformations (Table S1) 11

**S15.** Bibliographic summary of sequence context determinants affecting N-glycosylation efficiency (Table S2) 13

**S16.** DNA sequences of the constructs used in this study (Table S3) 14

**S17.** Multiple sequence alignment of the STT3A, STT3, AGLB1 and PGLB full protein sequences (Table S4) 17

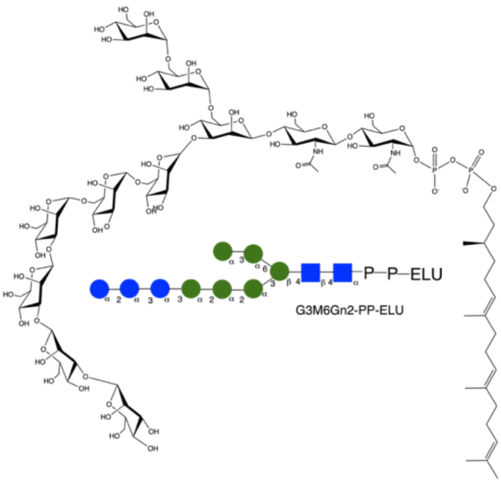

**Figure S1: Chemical and schematic (SNFG) representations of the lipid-linked oligosaccharide (LLO) used in MD simulations.** The oligosaccharide is covalently linked to a polyisoprenoid lipid chain via a pyrophosphate moiety. The oligosaccharide consists of two GlcNAc residues linked to a branched core of six mannoses, capped by three terminal glucose units. The anomeric center of the first sugar residue linked to the pyrophosphate is in the α configuration. The lipid moiety consists of a saturated isoprenyl unit bearing an *S* chiral center, followed by 2 trans-configured isoprenyl units and a terminal isoprenyl group. Abbreviations: Man, *D*-mannose; Glc, *D*-glucose; GlcNAc, *N*-acetyl-*D*-glucosamine.

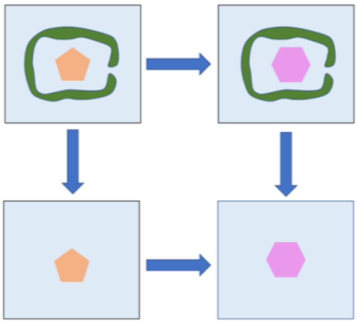

**Figure S2: Schematic representation of the thermodynamic cycle.** The alchemical transformations were performed along the horizontal pathways. In the upper row, the sequon mutation was carried out in the active site of OST-A embedded in a membrane and solvated in water, whereas in the second row the same sequon mutation was performed for the isolated peptide in water. Instead of directly calculating the relative binding free energy from the vertical transformations between the wild type (WT) and mutant states, the free energy difference was obtained by subtracting the free energies of the horizontal alchemical transformations, taking advantage of free energy being a state function. Orange and pink geometric shapes represent the WT and mutant sequons, respectively, while the green shape in the upper row represents the OST-A protein environment.

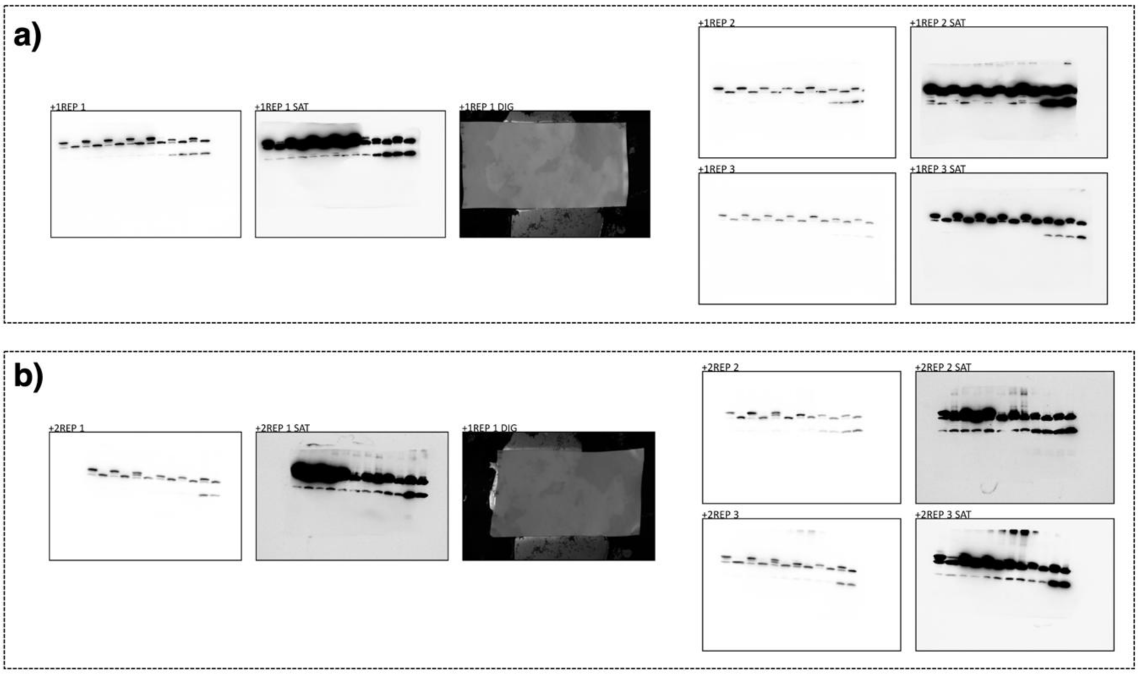

**Figure S3: Full Western blot membranes corresponding to Lep mutants at position +1 (A) and +2 (B).** "SAT" indicates saturated signal exposure. "REP" denotes independent biological replicates. "DIG" indicates a digital image showing the molecular weight ladder.

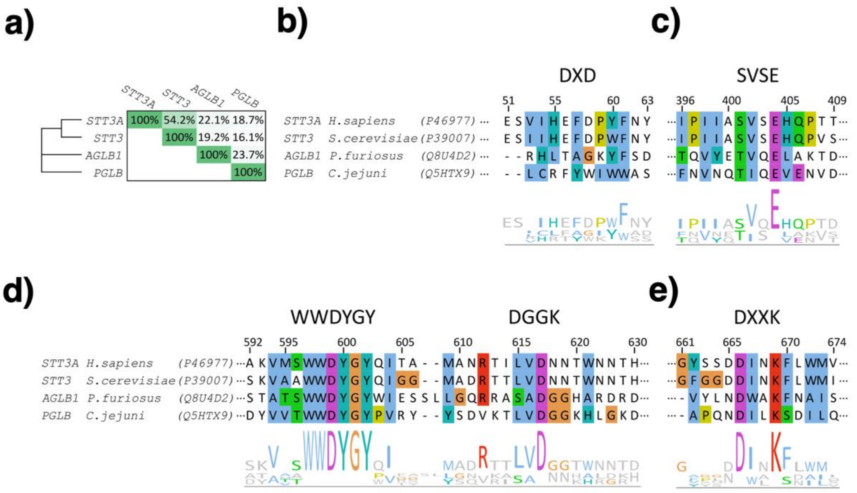

**Figure S4: Sequence conservation of the catalytic subunits of oligosaccharyltransferases from representative eukaryotic and prokaryotic organisms. (A)** Phylogenetic relationship and pairwise sequence identity among STT3A (*Homo sapiens*), STT3 (*Saccharomyces cerevisiae*), AGLB1 (*Pyrococcus furiosus*), and PGLB (*Campylobacter jejuni*). **(B,E)** Multiple sequence alignments of N-terminal domain motifs DXD and SVSE; and C-terminal domain motifs DGGK, WWDYG, and DK involved in the substrate binding and catalysis. Residues are coloured by ClustalX color scheme.

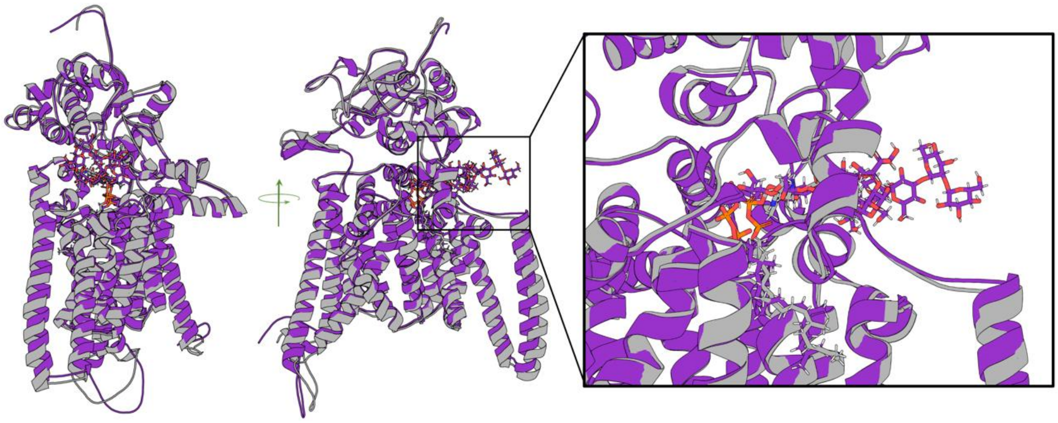

**Figure S5: Structural alignment of the model with the cryo-EM structure (PDB: 9N9J) of the OST complex.** Superposition of the enzyme model (cartoon representation in purple) with the cryo-EM structure of the OST complex (grey). The middle panel shows a rotated view of the structural overlap. The right panel provides a zoomed-in view of the active site region.

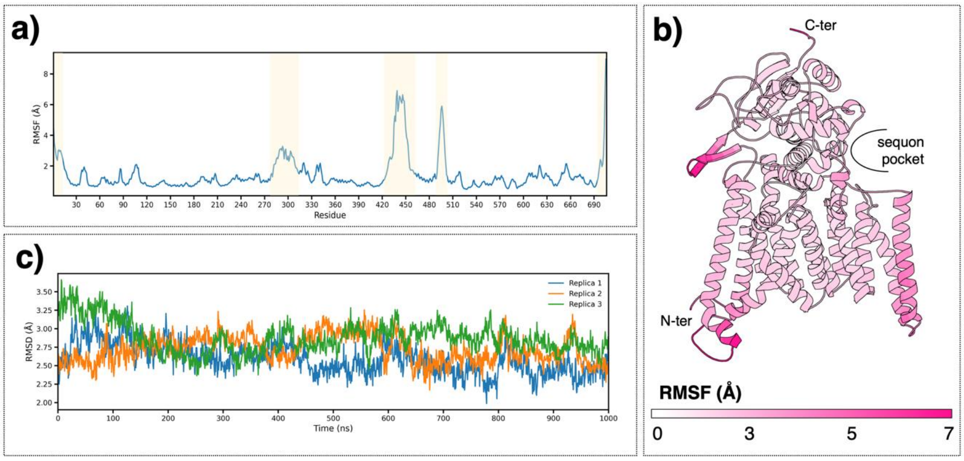

**Figure S6: Flexibility and stability of the OST-A model during molecular dynamics simulations. (A)** Average Cα root-mean square fluctuation (RMSF) values. Light yellow patches highlight regions of higher flexibility. **(B)** Structural representation of OST-A model with the residues colored by the RMSF value with the legend values. **(C)** Root-mean square deviations (RMSD) of the three independent replicas along the simulation time.

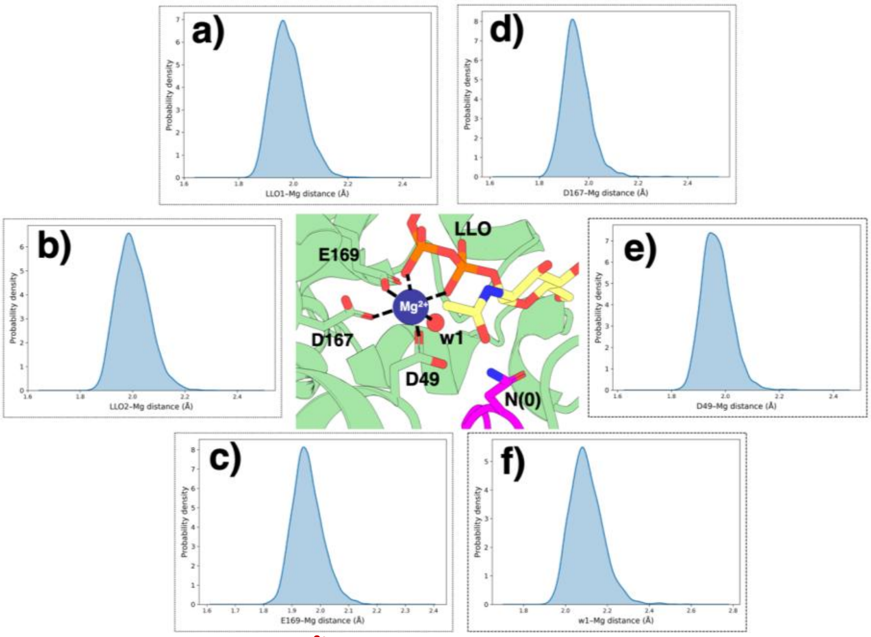

**Figure S7: Distribution of the distances between the $Mg^{2+}$ and the coordinating residues as depicted in the panel in the middle. (A)** LLO – O1; **(B)** LLO – O2; **(C)** E169; **(D)** D167; **(E)** D49; **(F)** Water.

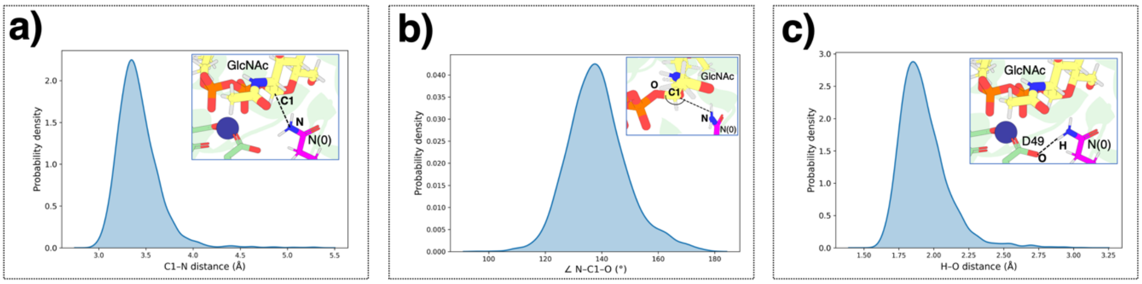

**Figure S8: Distributions of reaction-relevant geometries in the catalytic site during MD simulations. (A)** Distribution of values of the nucleophilic attack distance (N-C1) during the classical MD simulation; **(B)** Distribution of the N-C1-O angle during the MD simulation; **(C)** Distribution of values of the H atom of the amide group of the Asn(0) of the sequon and closest D49 oxygen atom (O-H) during the MD simulation.

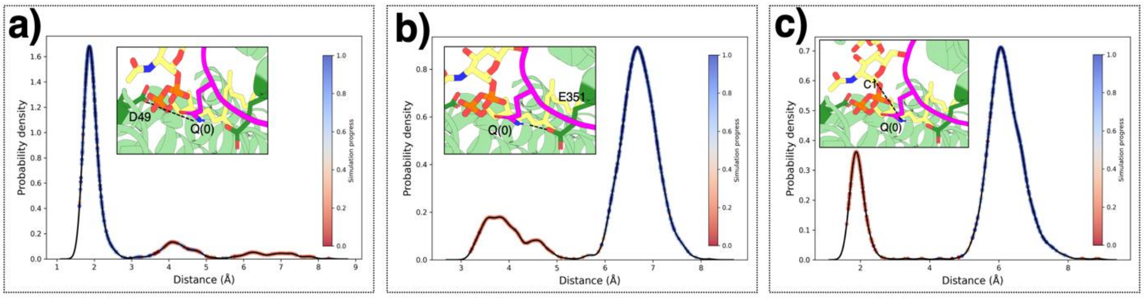

**Figure S9: Effects of the $N_0Q$ mutation on catalytically relevant interactions within the OST active site. (A)** Probability density distribution of the distance between D49 of the DXD motif and the side-chain amide group of the Q(0) of the sequon; **(B)** Probability density distribution of the distance between E351 of the SVSE motif and the side-chain amide group of the Q(0) of the sequon. In all panels, individual data points are colored according to simulation time to illustrate the irreversible shifts induced by the $N_0Q$ mutation. **(C)** Probability density distribution of the nucleophilic attack distance between the amide nitrogen of the Q(0) and the anomeric carbon of the sugar unit of the LLO substrate. Red corresponds to the beginning of the simulation and blue to the end of the simulation.

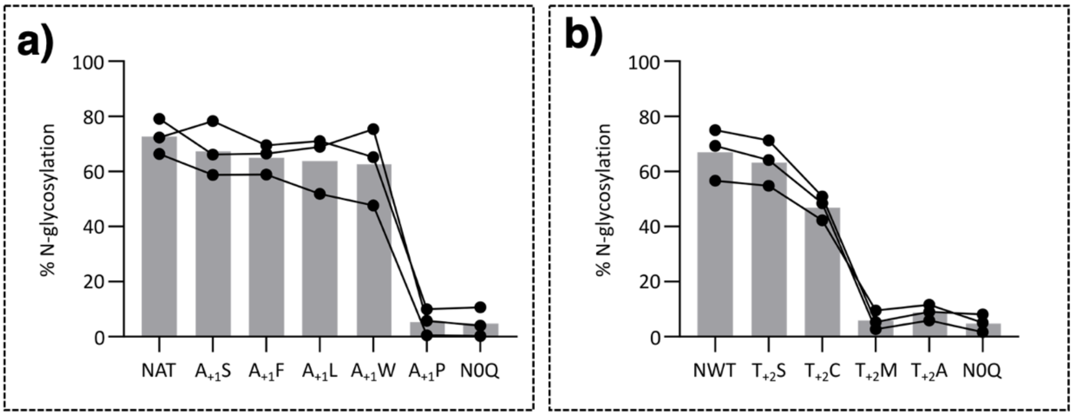

**Figure S10: Quantification of experimental N-glycosylation efficiency for the +1 (A) and +2 (B) positions.** N-glycosylation efficiencies were calculated as described for Figures 3e and 3g. Individual data points correspond to biological replicates, and lines connect measurements obtained from the same biological replicate across the different constructs to facilitate comparison of paired datasets.

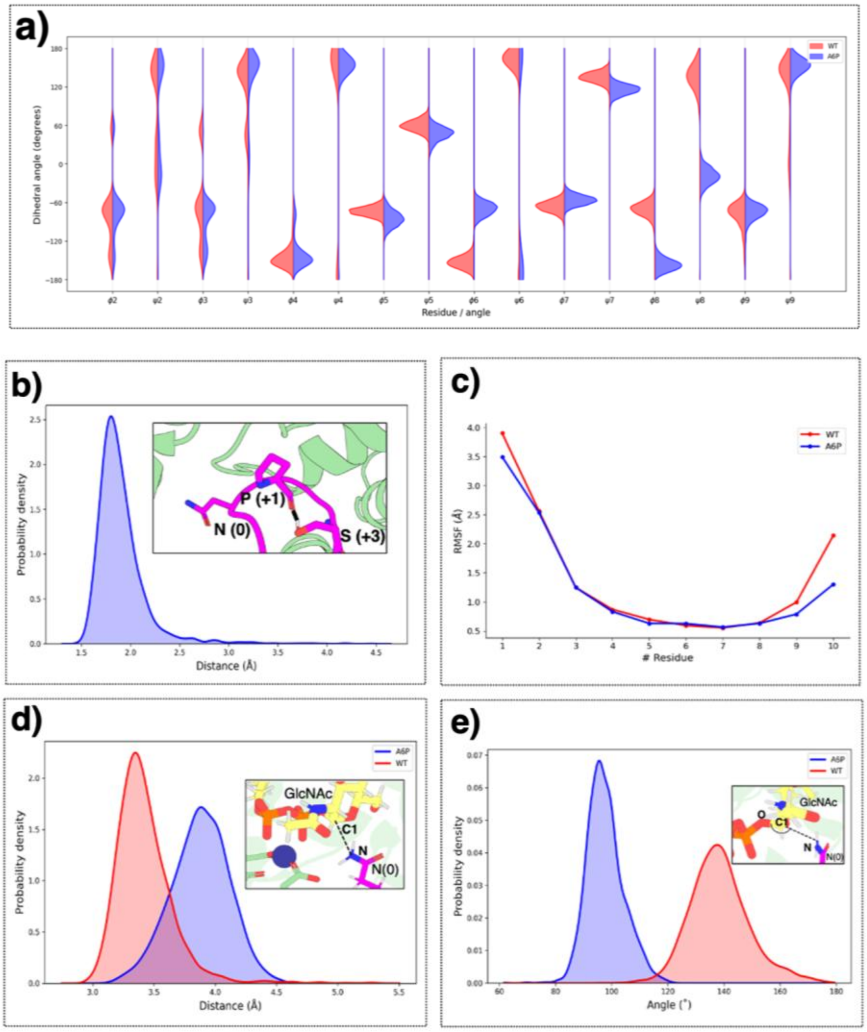

**Figure S11: Effect of the $A_{+1}P$ mutation on the sequon backbone conformation, flexibility and near-attack geometries. (A)** Violin plot representation of backbone φ and ψ dihedral angle distributions for residues across the WT (blue) and mutated $A_{+1}P$ sequon (red); **(B)** Distribution of the distance between sidechain hydroxyl group of Ser(+3) and backbone carbonyl of Pro mutant within the sequon. **(C)** Root-mean-square fluctuation (RMSF) of the sequon backbone in WT (red) and $A_{+1}P$ (blue). **(D)** The distribution of nucleophilic attack distances. **(E)** Distortion of the N–C1–O angle. WT distributions are shown in red and the $A_{+1}P$ mutant in blue.

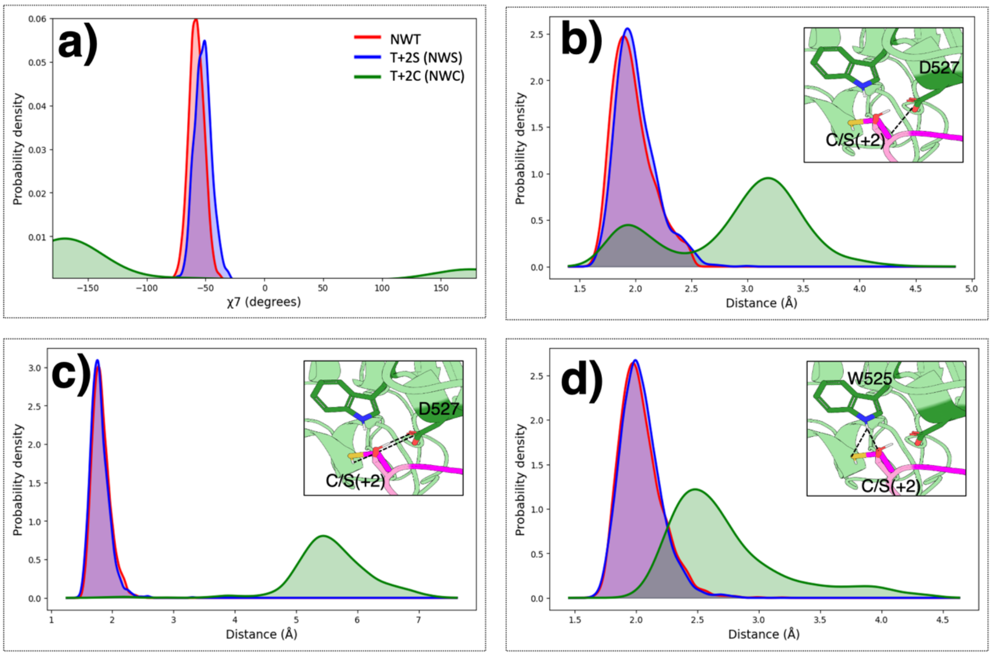

**Figure S12: Geometric determinants relevant for the +2 sequon recognition in the OST-A active site during MD simulations.** The common legend is given in the first panel. **(A)** Distribution of the χ-angle distributions of Ser/Thr hydroxyl and Cys thiol side chains at the +2 sequon position. **(B)** Distribution of the distance between the hydroxyl/thiol group of the +2 residue and the D527 carboxyl. **(C)** Distribution of the distance between the hydroxyl/thiol group of the +2 residue and the backbone amino group of the D527. **(D)** Distribution of the distance between the hydroxyl/thiol group of the +2 residue and the W525 side chain hydrogen.

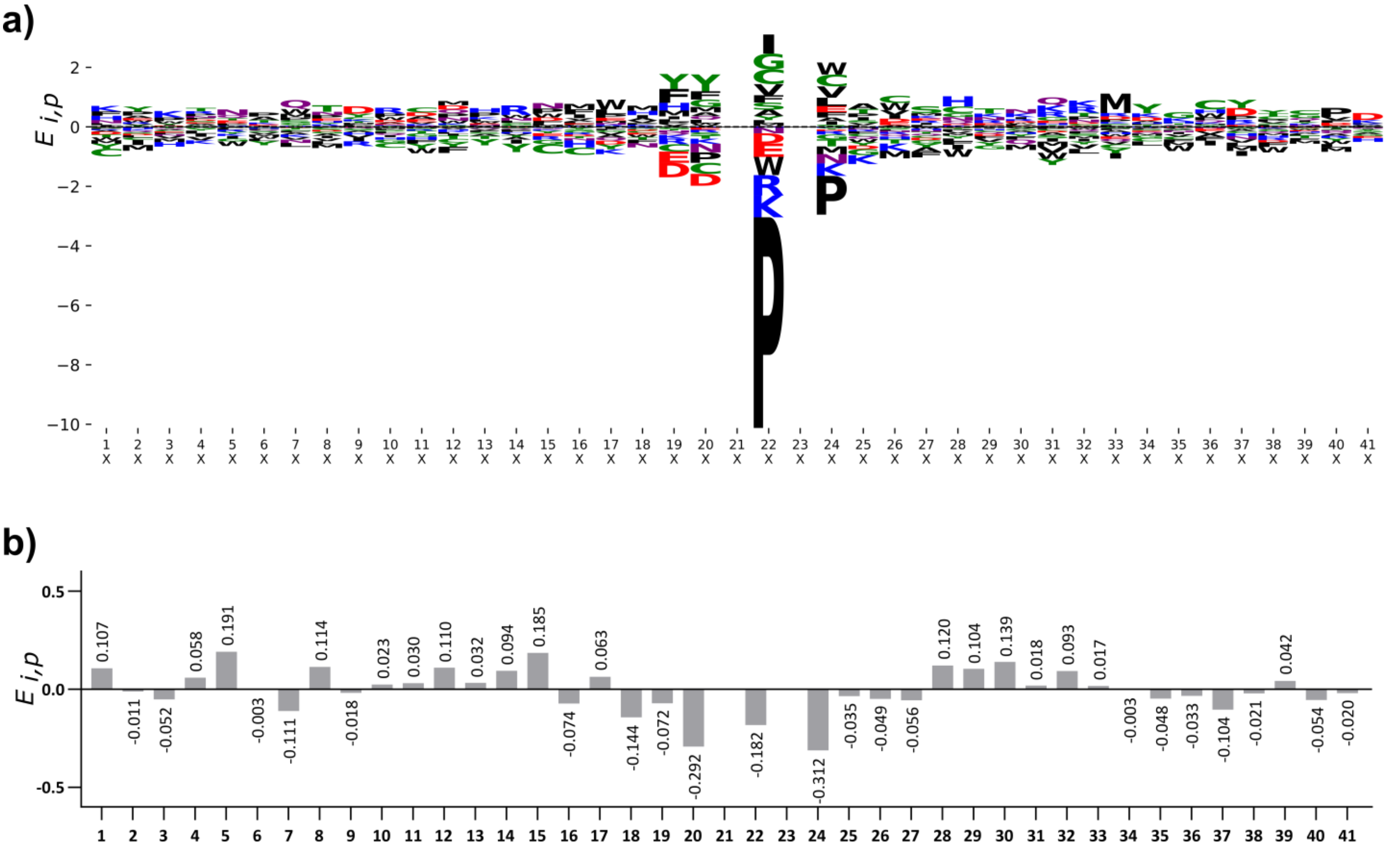

**Figure S13: Position-specific amino acid enrichment across the 41-residue glycosite window. (A)** Sequence logo showing the enrichment of all aminoacids at each position**. (B)** Position-specific enrichment of asparagine. Residues are colored by Chemistry (WebLogo3) color scheme. Polar-green (G, S, T, Y, C); neutral-purple (Q, N); basic-blue (K, R, H); acidic-red (D, E); and hydrophobic-black (A,V, L, I, P, W, F, M).

**Table S1: Free energy changes (kcal·mol⁻¹) associated with the alchemical transformations performed in the water and protein environment over five independent replicas.** The average values are given along with the standard deviations.

| **Transformation** | **$\Delta\Delta G = \Delta G_{prot} - \Delta G_{aq}$** | | | **2.85 ± 0.25** | | |
|---|---|---|---|---|---|---|
| **$N_0Q$** | environment | rep | $\Delta G_{aq}$ | environment | rep | $\Delta G_{prot}$ |
| | aqueous | 1 | 0.62 | protein | 1 | 3.85 |
| | | 2 | 0.60 | | 2 | 3.62 |
| | | 3 | 0.70 | | 3 | 3.53 |
| | | 4 | 0.73 | | 4 | 3.22 |
| | | 5 | 0.77 | | 5 | 3.44 |
| | mean | | 0.68 | mean | | 3.53 |
| | std | | 0.07 | std | | 0.25 |
| **Transformation** | **$\Delta\Delta G = \Delta G_{prot} - \Delta G_{aq}$** | | | **0.30 ± 0.14** | | |
| **$A_{+1}S$** | environment | rep | $\Delta G_{aq}$ | environment | rep | $\Delta G_{prot}$ |
| | aqueous | 1 | -10.21 | protein | 1 | -9.92 |
| | | 2 | -10.22 | | 2 | -9.88 |
| | | 3 | -10.22 | | 3 | -9.87 |
| | | 4 | -10.23 | | 4 | -10.18 |
| | | 5 | -10.24 | | 5 | -9.77 |
| | mean | | -10.22 | mean | | -9.92 |
| | std | | 0.01 | std | | 0.13 |
| **Transformation** | **$\Delta\Delta G = \Delta G_{prot} - \Delta G_{aq}$** | | | **0.69 ± 0.11** | | |
| **$A_{+1}F$** | environment | rep | $\Delta G_{aq}$ | environment | rep | $\Delta G_{prot}$ |
| | aqueous | 1 | -4.85 | protein | 1 | -4.20 |
| | | 2 | -4.86 | | 2 | -3.99 |
| | | 3 | -4.82 | | 3 | -4.24 |
| | | 4 | -4.87 | | 4 | -4.15 |
| | | 5 | -4.87 | | 5 | -4.28 |
| | mean | | -4.86 | mean | | -4.17 |
| | std | | 0.02 | std | | 0.10 |
| **Transformation** | **$\Delta\Delta G = \Delta G_{prot} - \Delta G_{aq}$** | | | **-0.84 ± 0.33** | | |
| **$A_{+1}W$** | environment | rep | $\Delta G_{aq}$ | environment | rep | $\Delta G_{prot}$ |
| | aqueous | 1 | -10.49 | protein | 1 | -11.00 |
| | | 2 | -10.43 | | 2 | -10.94 |
| | | 3 | -10.44 | | 3 | -11.53 |
| | | 4 | -10.40 | | 4 | -11.23 |
| | | 5 | -10.39 | | 5 | -11.67 |
| | mean | | -10.43 | mean | | -11.27 |
| | std | | 0.04 | std | | 0.29 |
| **Transformation** | **$\Delta\Delta G = \Delta G_{prot} - \Delta G_{aq}$** | | | **0.77 ± 0.36** | | |
| **$A_{+1}L$** | environment | rep | $\Delta G_{aq}$ | environment | rep | $\Delta G_{prot}$ |
| | aqueous | 1 | -0.83 | protein | 1 | 0.45 |
| | | 2 | -0.84 | | 2 | 0.18 |
| | | 3 | -0.78 | | 3 | -0.32 |
| | | 4 | -0.83 | | 4 | -0.47 |
| | | 5 | -0.77 | | 5 | -0.07 |
| | mean | | -0.81 | mean | | -0.04 |
| | std | | 0.03 | std | | 0.33 |
| **Transformation** | **$\Delta\Delta G = \Delta G_{prot} - \Delta G_{aq}$** | | | **2.50 ± 0.45** | | |
| | environment | rep | $\Delta G_{aq}$ | environment | rep | $\Delta G_{prot}$ |
| | aqueous | 1 | 19.32 | protein | 1 | 21.57 |
| | | 2 | 19.42 | | 2 | 22.10 |
| | | 3 | 19.74 | | 3 | 21.84 |

| | | | | | | |
|---|---|---|---|---|---|---|
| | | 4 | 19.00 | | 4 | 21.66 |
| | | 5 | 19.06 | | 5 | 21.88 |
| **$A_{+1}P$** | mean | | 19.31 | mean | | 21.81 |
| | std | | 0.27 | std | | 0.18 |
| **Transformation** | **$\Delta\Delta G = \Delta G_{prot} - \Delta G_{aq}$** | | | **2.43 ± 0.43** | | |
| | environment | rep | $\Delta G_{aq}$ | environment | rep | $\Delta G_{prot}$ |
| | aqueous | 1 | 0.99 | protein | 1 | 3.28 |
| | | 2 | 1.40 | | 2 | 3.49 |
| **$T_{+2}S$** | | 3 | 1.09 | | 3 | 3.18 |
| | | 4 | 1.05 | | 4 | 4.32 |
| | | 5 | 1.09 | | 5 | 3.54 |
| | mean | | 1.13 | mean | | 3.56 |
| | std | | 0.14 | std | | 0.40 |
| **Transformation** | **$\Delta\Delta G = \Delta G_{prot} - \Delta G_{aq}$** | | | **3.63 ± 0.19** | | |
| | environment | rep | $\Delta G_{aq}$ | environment | rep | $\Delta G_{prot}$ |
| | aqueous | 1 | 8.49 | protein | 1 | 12.12 |
| **$T_{+2}C$** | | 2 | 8.47 | | 2 | 12.32 |
| | | 3 | 8.46 | | 3 | 11.93 |
| | | 4 | 8.44 | | 4 | 11.90 |
| | | 5 | 8.54 | | 5 | 12.30 |
| | mean | | 8.48 | mean | | 12.11 |
| | std | | 0.04 | std | | 0.18 |

**Table S2: Bibliographic summary of sequence context determinants affecting N-glycosylation efficiency.** The acceptor asparagine (Asn) is shown in **red**. The residue highlighted in **bold** indicates the variable position examined in each study. Residues shown in regular font correspond to fixed positions explicitly defined in that paper. Amino acids associated with high or low N-glycosylation efficiency are summarized from the indicated references.

| Sequon context | Tested AA | High efficiency | Low efficiency | Ref. |
|---|---|---|---|---|
| ($^{Nt}$...-**X**-A-N-G-T-...$^{Ct}$) | F/H/W/Y | F/H/W/Y | -- | 1 |
| ($^{Nt}$...-**X**-K-N-G-T-...$^{Ct}$) | All | Y/M/C/W | R/K | 2 |
| ($^{Nt}$...-**X**-N-S-T-M-M-M-S-S-S-...$^{Ct}$) | All | F/S/L/G | P/M | 3 |
| ($^{Nt}$...-F-**X**-N-G-T-...$^{Ct}$) | All | C | W | 2 |
| ($^{Nt}$...-W-**X**-N-G-T-...$^{Ct}$) | All | C | -- | 2 |
| ($^{Nt}$...-R-**X**-N-G-T-...$^{Ct}$) | All | C/G | D | 2 |
| ($^{Nt}$...-N-**X**-S-...$^{Ct}$) | All | T/H/C/S | P/W/D/E | 4 |
| ($^{Nt}$...-N-**X**-T-...$^{Ct}$) | W/D/E/L/N/ G/R/H/S | W/D/E/L/N/G/R /H/S | -- | 5 |
| ($^{Nt}$...-N-**X**-S-...$^{Ct}$) | W/D/E/L/N/ G/R/H/S | G/R/H/S | W/D/E/L | 5 |
| ($^{Nt}$...-W-C-N-**X**-T-...$^{Ct}$) | All | G/Q/S/A | W/P/F/L/I/V | 2 |
| ($^{Nt}$...-R-D-N-**X**-T-...$^{Ct}$) | All | G/T/C/S/A | W/P/L/K | 2 |
| ($^{Nt}$...-N-L-**X**-...$^{Ct}$) | T/S/C | T/S | C | 6 |
| ($^{Nt}$...-N-L-T-**X**-...$^{Ct}$) | All | All (≠ P) | P | 7 |
| ($^{Nt}$...-N-L-S-**X**-...$^{Ct}$) | All | T/R/S/C | P/D/E/W/G | 7 |

**Table S3: DNA sequences of the constructs used in this study.** Start and stop codons are shown in bold. Sequences in lowercase denote vector-flanking sequences, while underlined sequences correspond to DNA encoding the transmembrane domains of Lep. Sequences shown in red encode N-glycosylation sequons, sequences shown in blue encode the c-Myc tag and sequences highlighted in grey encode the $^{Nt}$-A-Y-A-N-A-T-S-A-A-$^{Ct}$ peptide.

| DNA sequence for extended Lep variant in original pGEM vector: |
|---|
| gctgcaggtcgactctagagccacc**ATG**GCGAATTCCACCAGCCAGGGTTCTCAAC-<br>CGATCAACGCACAGGCGGCTCCGGTAGCGCAAGGTGGCAGTCAGGGAGAGTTTGCCCTGATTCTGGTGATTGCCACACTGGTGACGGGCATTTTAT<br>GGTGCGTGGATAAATTCTTTTTCGCACCTAAACGGCGGGAACGTCAGGCAGCGGCG-<br>CAGGCGGCTGCCGGGGACTCACTGGATAAAGCAACGTTGAAAAAGGTTGCGCCGAAGACTAGTTGGCTGGAAACCGGTGCTTCTGTTTTTCCGGT<br>ACTGGCTATCGTATTGATTGTGCGTTCGTTTATTTATGAACCGGTAC-<br>CGATCCCGTCAGGTTCGATGATGCCGACTCTGAACTCTACTGATTTTATTCTGGTAGAGAAGTTTGCTTATGGCATTAAAGATCCTATCTACCAGA<br>AAACGCTGATCGAAACCGGTCATCCGAAACGCGGCGATATCGTGGTCTTTAAATATCCGGAA-<br>GATCCAAAGCTTGATTACATCAAGCGCGCGGTGGGTTTACCGGGCGATAAAGTCACTTACGATCCGGTCTCAAAAGAGCTGACGATTCAACCGGG<br>ATGCAGTTCCGGCCAGGCGTGTGAAAACGCGCTGCCGGTCACCTACTCAAACGTGGAAC-<br>CGAGCGATTTCGTTCAGACCTTCTCACGCCGTAATGGTGGGGAAGCGACCAGCGGATTCTTTGAAGTGCCGAAACAGGAAACCAAAGAAAATGGA<br>ATTCGTCTTTCCGAGCGTAAAGAGACACTGGGTGATGTGACGCACCGCATTCTGACAG-<br>TGCCGATTGCGCAGGATCAGGTGGGGATGTATTACCAGCAGCCAGGGCAACAACTGGCAACCTGGATTGTTCCTCCGGGACAATACTTCATGATGG |

GCGACAACCGCGACAACAGCGCGGACAGCCGTTACTGGGGCTTTGTGCCGGAA-
GCGAATCTGGTCGGTCGGGCAACGGCTATCTGGATGAGCTTCGATAAGCAAGAAGGCGAATGGCCGACTGGTCTGCGCTTAAGTCGCATTGGCGG
CATCCAT**TAA**tagccatcttcgttcacgttt

DNA sequence for extended Lep variant in pHDM vector:

actttggcaaagaattccgcgggcggccgcgccacc**ATG**GCGAATTCCAC-
CAGCCAGGGTTCTCAACCGATCAACGCACAGGCGGCTCCGGTAGCGCAAGGTGGCAGTCAGGGAGAGTTTGCCCTGATTCTGGTGATTGCCACACT
GGTGACGGGCATTTTATGGTGCGTGGATAAATTCTTTTTCGCACCTAAACGGCGGGAAC-
GTCAGGCAGCGGCGCAGGCGGCTGCCGGGGACTCACTGGATAAAGCAACGTTGAAAAAGGTTGCGCCGAAGACTAGTTGGCTGGAAACCGGTGCT
TCTGTTTTTCCGGTACTGGCTATCGTATTGATTGTGCGTTCGTTTATTTATGAACCGGTAC-
CGATCCCGTCAGGTTCGATGATGCCGACTCTGAACTCTACTGATTTTATTCTGGTAGAGAAGTTTGCTTATGGCATTAAAGATCCTATCTACCAGA
AAACGCTGATCGAAACCGGTCATCCGAAACGCGGCGATATCGTGGTCTTTAAATATCCGGAA-
GATCCAAAGCTTGATTACATCAAGCGCGCGGTGGGTTTACCGGGCGATAAAGTCACTTACGATCCGGTCTCAAAAGAGCTGACGATTCAACCGGG
ATGCAGTTCCGGCCAGGCGTGTGAAAACGCGCTGCCGGTCACCTACTCAAACGTGGAAC-
CGAGCGATTTCGTTCAGACCTTCTCACGCCGTAATGGTGGGGAAGCGACCAGCGGATTCTTTGAAGTGCCGAAACAGGAAACCAAAGAAAATGGA
ATTCGTCTTTCCGAGCGTAAAGAGACACTGGGTGATGTGACGCACCGCATTCTGACAG-
TGCCGATTGCGCAGGATCAGGTGGGGATGTATTACCAGCAGCCAGGGCAACAACTGGCAACCTGGATTGTTCCTCCGGGACAATACTTCATGATGG
GCGACAACCGCGACAACAGCGCGGACAGCCGTTACTGGGGCTTTGTGCCGGAA-
GCGAATCTGGTCGGTCGGGCAACGGCTATCTGGATGAGCTTCGATAAGCAAGAAGGCGAATGGCCGACTGGTCTGCGCTTAAGTCGCATTGGCGG
CATCCATATGGAACAAAAACTCATCTCAGAAGAGGATCTG**TAA**ggatccaagcttatcgataccgtc

DNA sequence for modified Lep with the $^{Nt}$-A-Y-A-N-A-T-S-A-A-$^{Ct}$ peptide in pHDM vector:

actttggcaaagaattccgcgggcggccgcgccacc**ATG**GCGAATTCCAC-
CAGCCAGGGTTCTCAACCGATCAACGCACAGGCGGCTCCGGTAGCGCAAGGTGGCAGTCAGGGAGAGTTTGCCCTGATTCTGGTGATTGCCACACT
GGTGACGGGCATTTTATGGTGCGTGGATAAATTCTTTTTCGCACCTAAACGGCGGGAAC-
GTCAGGCAGCGGCGCAGGCGGCTGCCGGGGACTCACTGGATAAAGCAACGTTGAAAAAGGTTGCGCCGAAGACTAGTTGGCTGGAAACCGGTGCT
TCTGTTTTTCCGGTACTGGCTATCGTATTGATTGTGCGTTCGTTTATTTATGAACCGGTAC-
CGATCCCGTCAGGTTCGATGATGCCGACTCTGGCTTATGCCAACGCCACATCAGCCGCACTGGTAGAGAAGTTTGCTTATGGCATTAAAGATCCTA
TCTACCAGAAAACGCTGATCGAAACCGGTCATCCGAAACGCGGCGATATCGTGGTCTTTAAA-
TATCCGGAAGATCCAAAGCTTGATTACATCAAGCGCGCGGTGGGTTTACCGGGCGATAAAGTCACTTACGATCCGGTCTCAAAAGAGCTGACGAT
TCAACCGGGATGCAGTTCCGGCCAGGCGTGTGAAAACGCGCTGCCGGTCACCTACTCAAAC-
GTGGAACCGAGCGATTTCGTTCAGACCTTCTCACGCCGTAATGGTGGGGAAGCGACCAGCGGATTCTTTGAAGTGCCGAAACAGGAAACCAAAGA
AAATGGAATTCGTCTTTCCGAGCGTAAAGAGACACTGGGTGATGTGACGCACCG-
CATTCTGACAGTGCCGATTGCGCAGGATCAGGTGGGGATGTATTACCAGCAGCCAGGGCAACAACTGGCAACCTGGATTGTTCCTCCGGGACAATA
CTTCATGATGGGCGACAACCGCGACAACAGCGCGGACAGCCGTTACTGGGGCTTTGTGCCG-
GAAGCGAATCTGGTCGGTCGGGCAACGGCTATCTGGATGAGCTTCGATAAGCAAGAAGGCGAATGGCCGACTGGTCTGCGCTTAAGTCGCATTGG
CGGCATCCATATGGAACAAAAACTCATCTCAGAAGAGGATCTG**TAA**ggatccaagcttatcgataccgtc

Forward and reverse primers used for the construction of Lep mutants based on the modified Lep $^{Nt}$-A-Y-A-N-A-T-S-A-A-$^{Ct}$ plasmid:

| | |
|---|---|
| $A_{+1}S$ (NST) | CTTATGCCAACTCCACATCAGCCGCACTGGT<br>CAGTGCGGCTGATGTGGAGTTGGCATAAG |
| $A_{+1}F$ (NFT) | GGCTTATGCCAACTTCACATCAGCCGCACTGG<br>CAGTGCGGCTGATGTGAAGTTGGCATAAGCCAG |
| $A_{+1}L$ (NLT) | ACTCTGGCTTATGCCAACCTGACATCAGCCGCACTGGT<br>ACCAGTGCGGCTGATGTCAGGTTGGCATAAGCCAGAG |
| $A_{+1}P$ (NPT) | GGCTTATGCCAACCCCACATCAGCCGC<br>CGGCTGATGTGGGGTTGGCATAAGC |
| $N_0Q$ (QAT) | ACTCTGGCTTATGCCCAGGCCACATCAGCCGC<br>GCGGCTGATGTGGCCTGGGCATAAGCCAGAGT |

| | |
|---|---|
| $T_{+2}S$ (NWS) | GGCTTATGCCAACTGGTCCTCAGCCGCACTGG<br>CCAGTGCGGCTGAGGACCAGTTGGCATAAGC |
| $T_{+2}C$ (NWC) | CTGGCTTATGCCAACTGGTGCTCAGCCGCACTGGTAG<br>CTACCAGTGCGGCTGAGCACCAGTTGGCATAAGCC |
| $T_{+2}M$ (NWM) | CTGGCTTATGCCAACTGGATGTCAGCCGCACTGGT<br>ACCAGTGCGGCTGACATCCAGTTGGCATAAGCCAG |
| $T_{+2}A$ (NWA) | CTGGCTTATGCCAACTGGGCTTCAGCCGCACTGGT<br>ACCAGTGCGGCTGAAGCCCAGTTGGCATAAGCCAG |

**Table S4: Multiple sequence alignment of the STT3A, STT3, AGLB1 and PGLB full protein sequences.** The MSAs were generated using Clustal Omega**.**

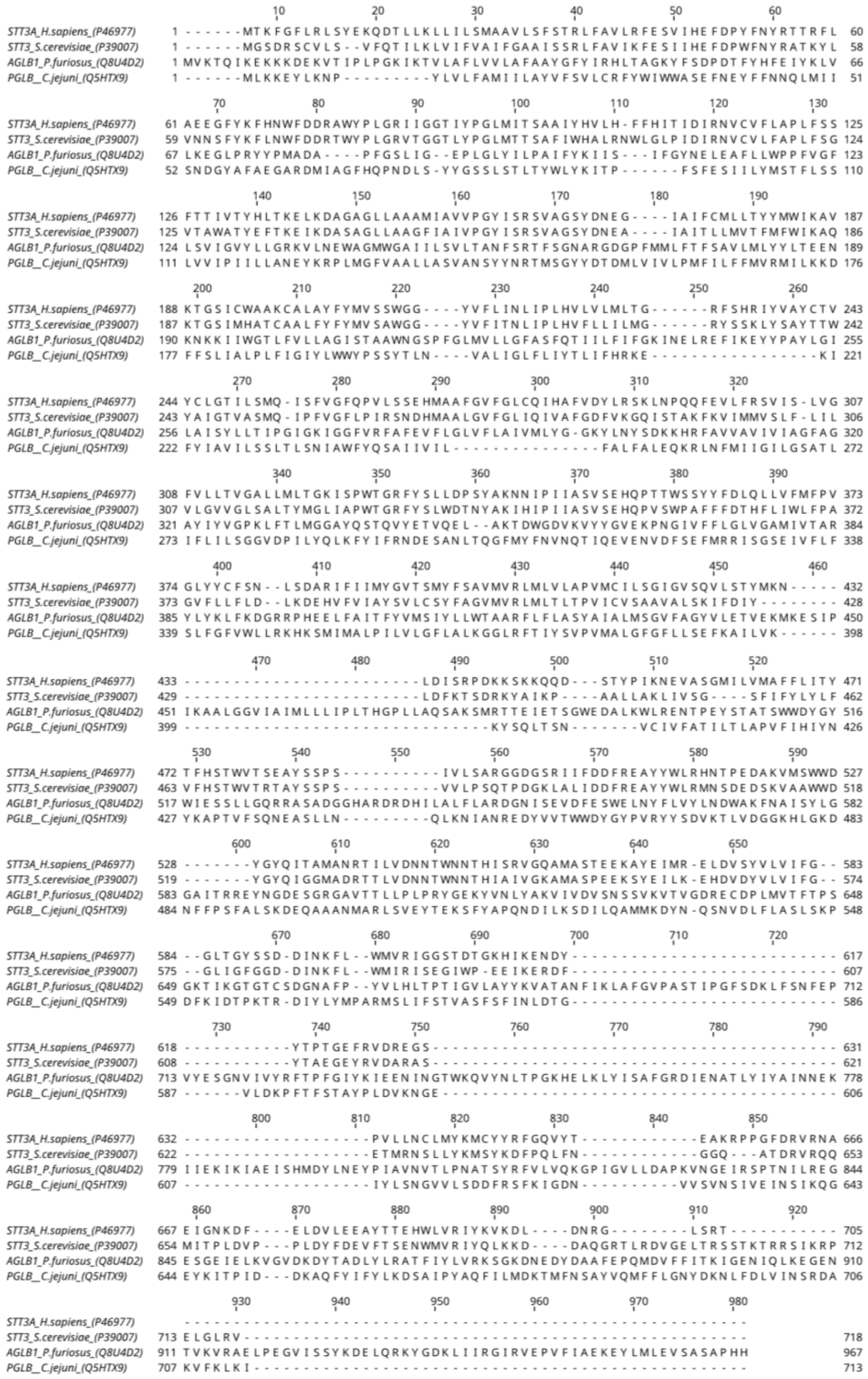
